# BESTDR: Bayesian quantification of mechanism-specific drug response in cell culture

**DOI:** 10.1101/2025.02.05.636681

**Authors:** Thomas O. McDonald, Simone Bruno, James P. Roney, Ioannis K. Zervantonakis, Franziska Michor

**Affiliations:** Department of Data Science, Dana-Farber Cancer Institute, Boston, MA, USA; Department of Biostatistics, Harvard T.H. Chan School of Public Health, Boston, MA, USA; Department of Stem Cell and Regenerative Biology, Harvard University, Cambridge, MA, USA; Center for Cancer Evolution, Dana-Farber Cancer Institute, Boston, MA, USA; Program in Computational and Systems Biology, Massachusetts Institute of Technology, Cambridge, MA, USA; Department of Bioengineering, UPMC Hillman Cancer Center, Swanson School of Engineering, University of Pittsburgh, Pittsburgh, PA, USA; The Eli and Edythe L. Broad Institute, Cambridge, MA, USA; Ludwig Center at Harvard, Harvard Medical School, Boston, MA, USA

## Abstract

Understanding drug responses at the cellular level is essential for elucidating mechanisms of action and advancing preclinical drug development. Traditional dose-response models rely on simplified metrics, limiting their ability to quantify parameters like cell division, death, and transition rates between cell states. To address these limitations, we developed *Bayesian Estimation of STochastic processes for Dose-Response* (BESTDR), a novel framework modeling cell growth and treatment response dynamics to estimate concentration-response relationships using longitudinal cell count data. BESTDR quantifies rates in multi-state systems across multiple cell lines using hierarchical modeling to support high-throughput screening. We validated BESTDR with synthetic and experimental datasets, demonstrating its robustness and accuracy in estimating drug response. By integrating mechanistic modeling of cytotoxic, cytostatic and other effects, BESTDR enhances dose-response studies, facilitating robust drug comparisons and mechanism-specific analyses. BESTDR offers a versatile tool for early-stage preclinical research, paving the way for drug discovery and informed experimental design.

## Main

In vitro cell culture experiments and dose-response modeling are fundamental to understanding drug response in pharmacology and serve an important function in the early phases of drug development^1^. These studies aim to quantify the relationship between drug concentration and cell growth using statistical models^2–4^. Dose-response modeling identifies efficacious drugs and, when combined with molecular profiling, helps uncover potential drug response biomarkers^5^.

In high-throughput screens, a range of cell lines are tested across panels of drugs to identify promising candidates for further study^6–9^. Data from such studies are often analyzed using real-time imaging technologies, which facilitate direct live-cell counting longitudinally during experiments^10,11^. These technologies also enable the labeling of distinct phenotypes, such as apoptotic cell states^12^, mutation status, epigenetic states^13,14^, and cell cycle phases^15^. The resulting multidimensional, longitudinal data provide an opportunity to better understand drug responses and cell kinetics, given appropriate statistical methods to leverage multiple cell states and time points simultaneously.

In traditional viability assay analysis, the relative fold-change in cell count is modeled as a function of drug concentration, and drug response is summarized using statistics for efficacy (e.g. E_max_) or potency (e.g. IC_50_)^16,17^. However, these classical models are sensitive to experimental conditions and do not account for sources of variability in the data which may stem from observation error, varying experimental conditions, or intrinsic stochasticity in cell cycling times. To address these challenges, time-invariant metrics, such as the Growth Rate (GR) approach, were developed that are not sensitive to experimental duration^18,19^. However, these metrics do not differentiate between mechanisms of action such as cytostatic (i.e. growth inhibition) and cytotoxic (i.e. cell killing) effects^19,20^. A similar approach models viable cell counts to estimate the cytotoxic and cytostatic responses with Gaussian process concentration-response curve for each response^21^. These methods are restricted to single cell states, limiting their usefulness for modeling complex mechanisms such as cell cycle transitions or cell state changes. These limitations impede the investigation of distinct drug mechanisms targeting specific states.

Here we present a new framework, *Bayesian Estimation of STochastic processes for Dose-Response* (BESTDR), for measuring the drug response of cell-kinetic mechanisms in cell culture experiments. Using longitudinal cell count data under varying drug concentrations (Fig. 1A), we model cell population growth as a branching process^22–24^. This approach describes population growth based on the dynamics of individual cells that may undergo fates such as division or death (Fig. 1B). Each of these fates can be expressed as functions of drug concentration (Fig. 1C). The goal of BESTDR is to use data and model specifications (Fig. 1A,B) to estimate the concentration-response function for each rate, corresponding to a specific fate or transition event (Fig. 1C). BESTDR estimates the posterior distribution of each parameter from which concentration-response curves can be constructed (Fig. 1D). These concentration-response estimates then provide insight into the drug’s mechanism of action. For example, the dynamics of cell division and death are different for cytostatic and cytotoxic drugs, and drugs may act according to either mechanism, or a combination of the two, even if the net growth dynamics are the same (Fig 1E).

**Figure 1.**
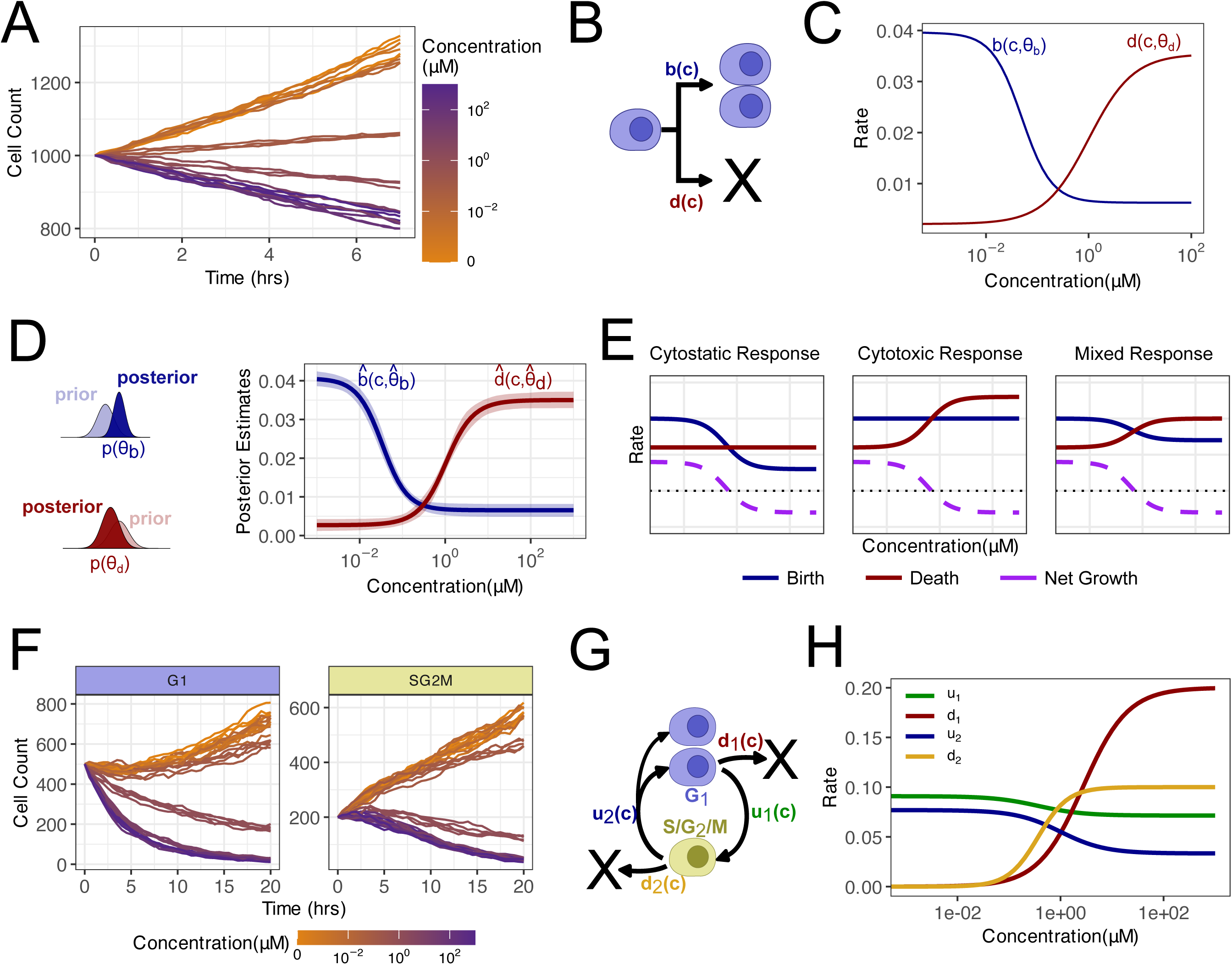
BESTDR workflow uses viable cell counting data to inform a branching process model of cell growth in order to estimate concentration-response parameters. **A** Longitudinal cell counting data is used to model the effects of drug concentration on cell growth in viable cells where cell growth is observed in culture in the presence of varying drug concentrations **B** Branching process models are used to represent cell state transitions of individual cells fates where each transition rate represents the frequency of that event occurring in order to describe the population growth. **C** Transition rates can be described by a concentration-response function parameterized by unknown parameters that BESTDR attempts to estimate in order to estimate the function. **D** BESTDR uses the model assumptions along with priors to update the normal likelihood model for cell counts, estimating each of the parameters of the concentration-response curve. The resulting concentration-response curve is a posterior distribution built on the posterior parameter estimates. **E** Similar net growth curves (purple) can come from different birth and death rates which represent mechanisms of response to a drug. Cytostatic and cytotoxic responses are represented by changes in the birth and death rates, respectively, while a mixed response can also arise. BESTDR is able to distinguish these responses by estimating the separate curves. **F** Cell-state specific phenotypes allow counting cells in individual states such as G1 or S/G2/M provides more dimensions of data to estimate specific drug effects. **G** Multitype branching process models can define additional cell specific events like cell phase transitions or phase-specific death. **H** BESTDR includes multitype methods in the same framework to estimate state-specific transition rates and further understand how cell lines respond to drugs with respect to specific mechanisms.

The BESTDR framework can also utilize data from multiple cell states to estimate concentration-response relationships of complex systems, such as transitions between states or the cell cycle (Fig 1F, G). This capability allows for the estimation of drug-specific responses on various mechanisms of cell growth, death and state switching without requiring additional mathematical complexity (Fig. 1H). Additionally, BESTDR supports high-throughput screening data by leveraging hierarchical modeling to account for experimental variability across different experiments or cell lines. By democratizing the modeling of more complex systems and accommodating variability inherent in large datasets, BESTDR becomes accessible for diverse applications, with the code available as an R package on GitHub^25^. In sum, BESTDR addresses a gap in modeling cell population dynamics in terms of the underlying mechanisms of drug response; such an approach is essential for investigating the effects of drugs on cell kinetics and serves as an important step in early drug development^15,26^.

## Results

### BESTDR is a statistical framework for estimation of drug response

Standard viability assays (Fig. 1A) measure drug response by normalizing viable cell counts at each concentration and time point to levels of control experiments and fitting a curve to this data to estimate parameters like the half-maximal inhibitory concentration (IC_50_)^27,28^. To overcome limitations of existing methods due to experimental conditions, noise, and a lack of mechanistic interpretation of drug response, we developed the BESTDR framework for systematically estimating drug response from viability assays. BESTDR models the counts of different cell states over time using a multitype continuous-time Markov branching process (CTMBP, Fig. 1B). In this stochastic framework, individual cells undergo fate events—such as cell division, death, or transitions between cell states—independently, with the time between events following an exponential distribution^22^. This approach is appropriate for modeling cell populations because it captures the inherent randomness in cell fate decisions at the single-cell level.

Each event has an associated rate parameter that can be described by a concentration-response (CR) function (Fig. 1C). BESTDR estimates the parameters of the CR functions for each rate and allows for prediction of the concentration-response curve and cell growth trajectories (Fig. 1D). To estimate the parameters of the CR functions, BESTDR employs the Hamiltonian Monte Carlo method^29,30^, a type of Markov Chain Monte Carlo algorithm that efficiently samples from posterior distributions. This estimation process uses a multivariate normal likelihood function to model observed cell counts over time which only requires numerically computing the mean and variance at each concentration and time (Methods)^23,31^. This approximation reduces computational complexity and facilitates efficient parameter estimation. Computing the variance of the branching process allows attributing variability to differences in transition dynamics, allowing us to estimate additional mechanistic parameters using the same data. Furthermore, BESTDR accounts for measurement errors from instrument miscounting by including a constant error term in the variance, maintaining accuracy (Methods, SI). Our method thus represents a statistically rigorous and efficient approach to determine parameters of mechanistic models of drug response from standard cell assay datasets.

### Deconvolving Cytostatic and Cytotoxic Drug Responses in silico

To validate BESTDR, we conducted in silico simulations of cell growth and death during treatment with 12 different drug concentrations. In this approach, each cell either divides into two daughter cells with rate b(c) or dies with rate d(c) where c is the drug concentration. Changes in b(c) or d(c) correspond to cytostatic and cytotoxic effects of the drug, respectively. Note that changes in either rate, or both, can result in the same net growth curve when the birth and death rates offset each other (Fig. 1E). Our approach deconvolves the estimates for birth and death rates separately using the number of viable cells which serves as the input into BESTDR; the net growth rate is then defined as the difference between birth and death rates. To account for technical variability, we introduced a 3% error to the observed cell counts, representing cell segmentation errors between observed and true cell counts that is similar to the accuracy of modern segmentation methods^32–34^.

We simulated data by first selecting parameters for the four-parameter logistic concentration-response curves for the birth and death rates (Ext. Data Table 1), which were then used to calculate the birth and death rates at each concentration and to determine simulated cell counts over time (Fig. 2A). The simulations were run for 72 hours with observations recorded every 4 hours to match standard experimental protocols. Using all time points, we then estimated the birth and death rates at each concentration independently to assess whether BESTDR could accurately recapitulate cell growth dynamics from the simulated data. At each concentration, the true birth and death rates fell within the 90% credible intervals (Fig. S1A, Ext. Data Table 2). We then used BESTDR to estimate the concentration-response (CR) curves using cell counts from all concentrations. All parameters of the CR curves fell within the 90% credible intervals of the posterior distributions (Fig S1B, Ext. Data Table 1). The largest relative errors between the true values and the posterior means were observed for the b_50_ (the concentration at which the birth rate is halfway between both asymptotes) and the Hill coefficient for the birth rate, b_h_, with relative errors of approximately 12.7% and 14%, respectively (Fig. S1B, Ext. Data Table 1). Importantly, the 90% credible bands for both the birth and death rates encompassed the true parameter values across all concentrations (Fig. 2B). This finding demonstrates that BESTDR can accurately estimate dynamic rates and provides mechanism-specific insights about cell division and death beyond net growth.

**Figure 2.**
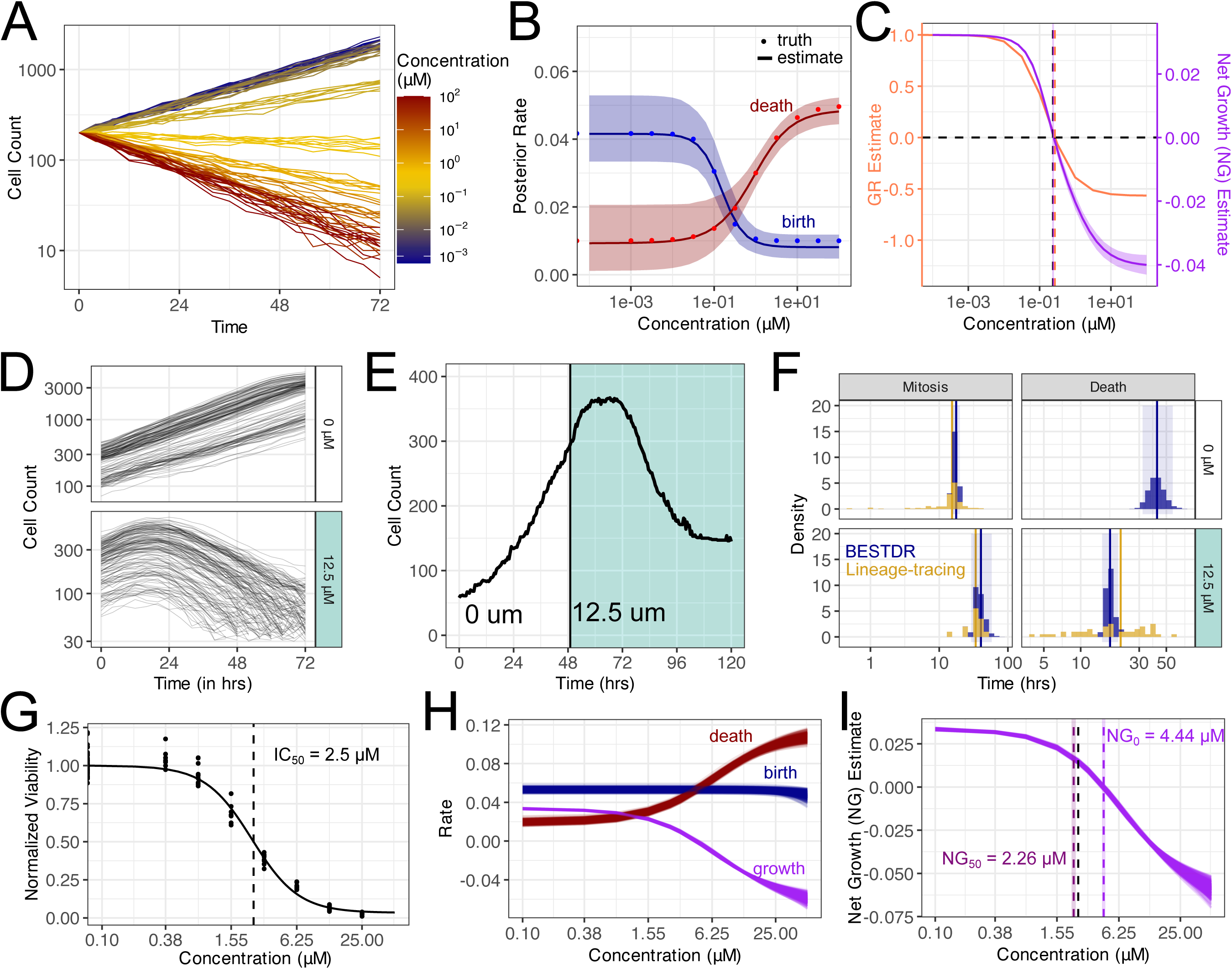
BESTDR estimation can deconvolve birth and death rates in viable cell counting experiments. **A** Simulation of longitudinal cell growth data following a four-parameter concentration response curve at various concentrations. **B** Posterior mean and 90% credible bands for the birth and death concentration-response curves built from the posterior distributions for the parameters accurately recapitulate the true concentration response. **C** Comparison between the Net Growth (purple) and Growth Rate (orange) methods show similar predicted concentrations at which cell turnover is 0. However, the total efficacy is different since Net Growth is in hr^-1^ units and provides a measure of the total change in cycling rate and Growth Rate is unitless as a measure of relative response. **D** HCT-116 p53-VKI cells were seeded and grown in cell culture for 96 hours in control (white) and 12.5 μM cisplatin (turquoise), recording viable cell count every 4 hours (dataset 1). **E** Live-cell lineage tracing experiments of HCT-116 p53-VKI cells (dataset 2) recording all cell fates have similar growth and response to cisplatin after treatment (turquoise) which is used to validate mitosis and death rates estimated from BESTDR. **F** BESTDR estimates of the average time between mitosis and death events using only cell counts (blue) is compared to the distribution of mitosis and death times in the lineage tracing data (gold) showing agreement in the estimates. **G** Normalized viability estimates of a concentration response and the corresponding IC_50_. **H** The birth (blue) and death (red) response curves show cisplatin is primarily cytotoxic with little to no effect on cycling time. **I** The net growth rate (purple) concentration-response describes the change in cell turnover with increasing concentration. The NG_50_ is similar in value to the 72-hour IC_50_, but the NG_0_ is more informative as the concentration where no net growth occurs.

We then calculated the net growth rate as the difference of the estimated birth and death rates and compared it to the GR curve^19^ derived from the same data (Fig. 2C). The average concentration at which the posterior net growth rate is 0 (NG_0_) is 0.251 μM (90% CI: [0.226, 0.275] μM), closely matching the estimated GR_0_ of 0.264 μM as expected since the NG_0_ and GR_0_ measure the same theoretical value. Our findings indicate that BESTDR accurately recapitulates summary statistics from in silico experiments while offering estimates of cell growth and death rates and drug efficacy in the original units of measurement (hr^-1^) as opposed to a relative response which is unitless as in the GR.

### BESTDR predicts in vitro cisplatin birth and death concentration-response curves

To validate BESTDR with experimental data, we compared BESTDR estimates obtained from cell counting experiments over multiple replicates to observed mitosis and death times in single cell live-cell lineage-tracing experiments. To obtain the first dataset, HCT-116 p53-VKI cells were treated with cisplatin and viable cell counts were recorded every 4 hours for 72 hours, both without drug and with 12.5 μM cisplatin, for multiple replicates (Methods). Cells grew exponentially in control and cisplatin, but started undergoing apoptosis approximately 20 hours after treatment initiation (Fig. 2D). To obtain the second dataset^35,36^, HCT-116 cells were cultured without cisplatin for 2 days followed by cisplatin treatment at 12.5 μM for 3 days; throughout the entire duration of the experiment, each cell’s fates were tracked and recorded (Fig. S1E). The two datasets are comparable in terms of the total number of viable cells at each timepoint, with the second dataset providing detailed information on cell fates that can be used to validate estimates obtained using BESTDR applied to the first dataset. We observed doubling times of 20.3 hours in dataset 1 vs. 20.0 hours in dataset 2 in control conditions, and similar maximum cell counts at around 16 to 20 hours post-treatment before the drug effect manifested in dataset 2, suggesting that the different methods of tracking cell growth yield comparable results (Fig. 2D,E).

Using BESTDR, we then estimated the birth and death rates from the count data in dataset 1 and compared them to the time-to-mitosis and time-to-death distributions from dataset 2. The estimated mean time to mitosis was 17.6 hours (95% CI: [15.7,19.9]) for control cells and 40.4 hours (95% CI: [29.2,58.0]) for treated cells, closely matching the lineage-tracing means of 15.4 and 34.0 hours, respectively (Fig. 2F). The estimated mean times to death under cisplatin were 42.0 hours (95% CI: [32.0,64.1]) for control and 17.4 hours (95% CI: [15.0,21.4]) for treated cells, compared to 21.2 hours observed in the lineage tracing data. Note that the distribution of dead cells in the control setting cannot be estimated, likely due to the low cell count and short tracking time (Fig 2F). In sum, our observations demonstrate that BESTDR provides estimates comparable to high-resolution cell-fate tracking without the need for extensive individual cell monitoring.

To investigate BESTDR’s ability to estimate a concentration-response curve, we then treated HCT-116 p53-VKI cells with eight concentrations of cisplatin and tracked viable cell counts for 72 hours (Fig. S1D). Due to the 20-hour delay in cisplatin’s effect, we analyzed data after 25 hours for higher concentrations to focus on the exponential growth phase. We chose a 3-parameter logistic function to model the 72-hour viability curves, yielding an estimated IC_50_ of 2.5 μM (Fig. 2G). BESTDR revealed that the death rate increased at lower concentrations while the birth rate remained largely constant, changing by only about 0.001 hr^⁻¹^ across the tested range. This observation suggests that cisplatin primarily induces apoptosis rather than inhibiting cell division, with a net change in death rate of 0.080 hr^⁻¹^ (Fig. 2H). This finding aligns with previous studies^37^ and the lineage-tracing data showing similar mitotic times but differing death rates between untreated and cisplatin-treated cells (Fig. S1C).

We used BESTDR to estimate the concentration at which the net growth rate is half that of the control (NG_50_) and 0 (NG_0_), respectively. The NG_50_ was estimated as 2.26 μM (90% CI: [2.16,2.37] μM, Fig. 2I) comparable but lower than the IC_50_ of 2.5 μM (Fig 2G), while the NG_0_ was estimated as 4.44 μM (90% CI: [4.28,4.60] μM, Fig. 2I). By revealing that cisplatin predominantly increases the death rate while keeping the cell birth rate largely unaffected, BESTDR offers valuable mechanistic insights into drug action.

### Accounting for Cell Clearance in Multi-type Models

We next investigated whether incorporating both viable and dead cell counts improves treatment response estimation compared to using viable counts alone. To this end, we analyzed in silico data obtained in a setting where viable cells may divide or die with logistic concentration-response functions, b(c) and d(c), and dead cells clear from the population at a constant rate, k (Fig. 3A,B). Clearance represents loss of adherence to the plate when cells are no longer counted and appear as a decrease in dead cell counts^38^. We simulated cell growth during treatment at 10 different concentrations, tracking viable and dead cells with clearance set to zero (Fig. 3C) and 0.02 hr^-1^ (Fig. 3D) and added random noise (Ext. Data Table 3). This data was then used to estimate birth and death rates using (1) BESTDR applied to viable counts (“Live BESTDR”), (2) BESTDR applied to viable and dead counts, accounting for dead-cell clearance (“Live-dead BESTDR”), and (3) the Static/Toxic GR Ordinary Differential Equation (“ODE”) model^26^, which estimates rates from viable and dead counts without accounting for clearance. When clearance was zero, all models closely fit the true birth and death rate curves. Live BESTDR also provided accurate estimates, with area between the curves (ABC) values of 0.023 (birth rate) and 0.029 (death rate), although with slightly wider credible intervals due to the use of only viable cell data (Fig. 3E, Ext. Data Table 3). Live-dead BESTDR achieved similar fits, with ABC values of 0.005 (birth rate) and 0.001 (death rate), and 95% credible intervals encompassing the true curves (Fig 3F). The ODE model had an ABC of 0.003 for the birth rate and 0.010 for the death rate, indicating high accuracy (Fig. 3G). These results suggest that live cell counting alone is sufficient for estimating birth and death rates when there is no dead cell clearance.

**Figure 3.**
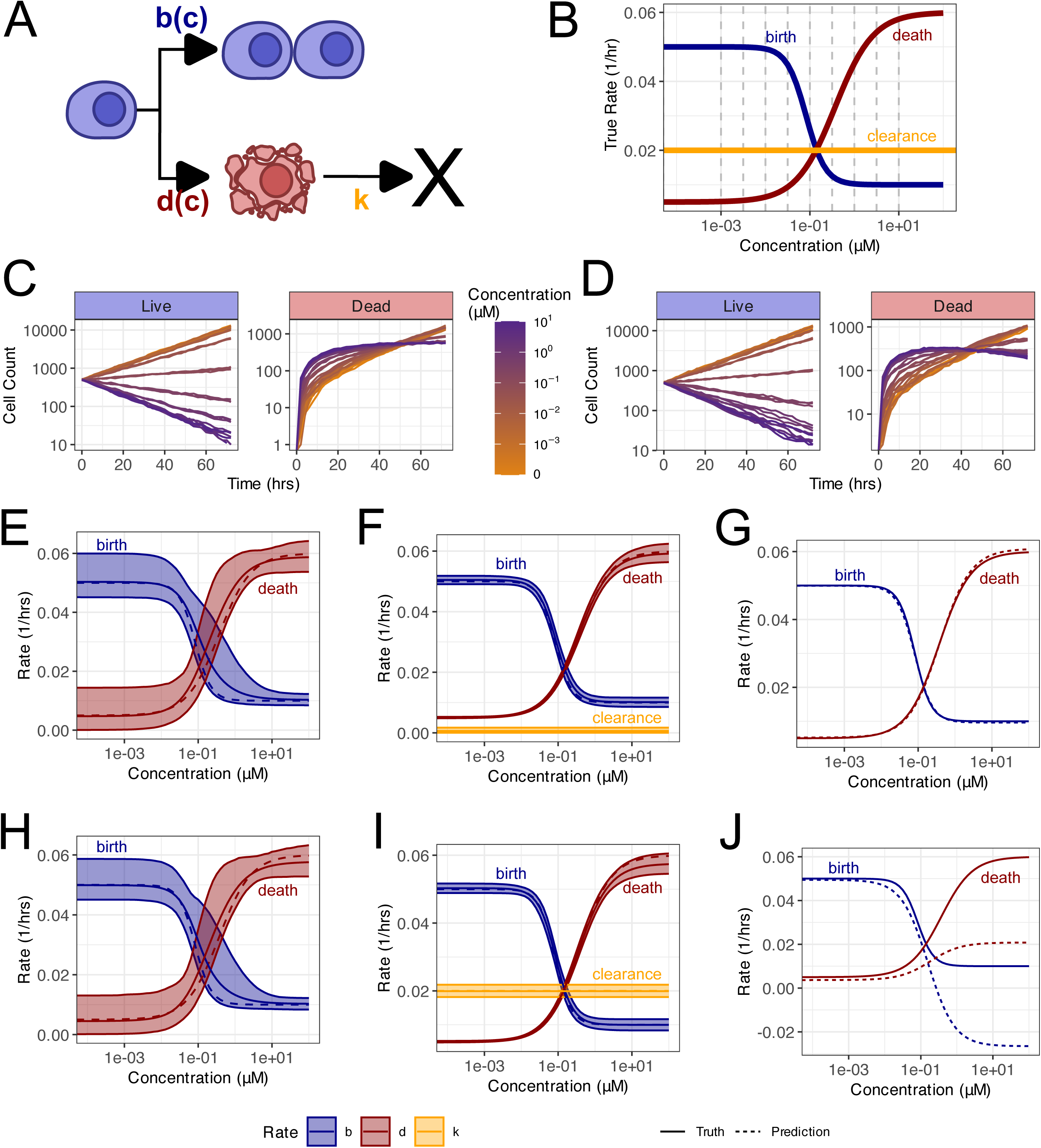
Comparing estimation of birth and death rates using viable and dead cell counts. **A** A model for cell growth and dead cell clearance. Viable cells can split or die with concentration-dependent rate functions b(c) and d(c) respectively while dead cells clear with constant rate k independent of drug concentration. **B** The concentration-response for the birth and death rates are modeled as 4-parameter logistic functions and constant clearance. **C** Simulations of trajectories from a model with birth and death rates with no clearance (k=0 hr^-1^) counting both viable and dead cells shows a monotonically increasing dead cell count. **D** A scenario with clearance (k=0.02 hr^-1^) allows dead cell counts to decrease at higher concentrations when the pool of viable cells also decreases. **E** Live-only BESTDR estimates and 90% credible bands of the concentration response curves with data where no clearance occurs. **F** Live-dead BESTDR estimates and 90% credible bands of the concentration response curves with data where no clearance occurs. **G** Static-toxic GR estimates of the concentration-response curves with data where no clearance occurs. **H** Live-only BESTDR estimates and 90% credible bands of the concentration response curves with data where the true clearance rate is 0.02 hr^-1^. **I** Live-dead BESTDR estimates and 90% credible bands of the concentration response curves with data where the true clearance rate is 0.02 hr^-1^. **J** Static-toxic GR estimates of the concentration-response curves with data where the true clearance rate is 0.02 hr^-1^.

When clearance was included, Live BESTDR maintained similar levels of accuracy to the no-clearance scenario, with ABC values of 0.025 (birth rate) and 0.028 (death rate) (Fig. 3H). Live-dead BESTDR accurately recapitulated the original rates, with ABC values of 0.002 (birth rate) and 0.005 (death rate), and the true dose-response curves falling within the 95% credible intervals (Fig. 3I). In contrast, the ODE model’s estimates deviated significantly from the true values at higher concentrations and even produced negative birth rates (Fig. 3J), with ABC values increasing to 0.126 (birth rate) and 0.128 (death rate).

Despite a slight increase in standard error of live-only BESTDR relative to live-dead BESTDR, all parameters in the BESTDR simulations fell within the 95% credible intervals (Ext. Data Table 3). These results illustrate that BESTDR models can effectively estimate birth and death rates even when dead-cell clearance is unknown, outperforming ODE models that fail to account for clearance. These observations highlight BESTDR’s robustness and practicality for analyzing cell growth data without the need for additional measurements of dead cells.

### Hierarchical Modeling for High-Throughput Drug Screens

We then extended the live-only BESTDR model to analyze high-throughput drug screens involving multiple cell lines. To account for cell line-specific diversity in drug responses^4,9,17^, we modeled drug response of all cell lines in a hierarchical framework, which assumes that individual curves are related^39^. Each cell line follows a birth-death process, as before (Fig. 4A), but the rate parameters are now realized from shared probability distributions, reflecting common dynamics and drug responses across cell lines, resulting in cell-line specific curves for each rate (Fig. 4B). We applied this hierarchical model to response data from eight breast cancer cell lines treated with 25 drugs over concentrations ranging from 1.8 x 10^-5^ μM to 3.9 μM^40^. Each drug was modeled with BESTDR independently, estimating cell line-specific parameters and hyperparameters specifying the shared distributions.

**Figure 4.**
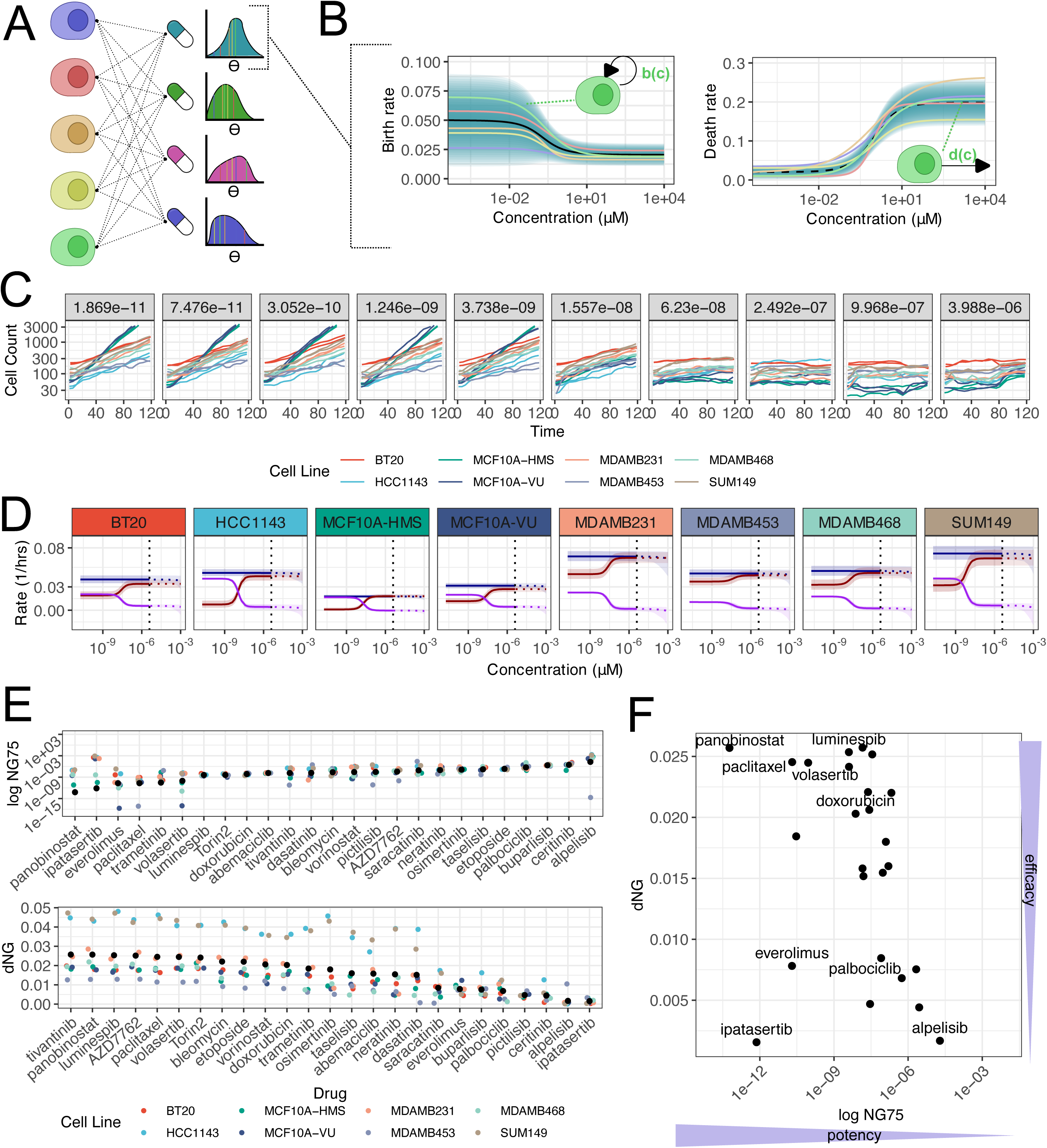
Hierarchical mechanistic concentration-response models use data from multiple cell lines for more robust estimates. **A** High throughput drug screens collect viability data across combinations of multiple cell lines for multiple drugs. The parameters that form the concentration response curve come from a distribution parameterized by hyperparameters. We model each cell line’s birth and death rate parameters as realizations from distributions such that there is some similarity across cell lines of the same type. **B** An individual cell line has birth and death concentration-response curves that are realizations from a distribution of possible curves that is built from the hyperparameters for the parameters that describe each curve. **C** Longitudinal cell counts over 8 breast cancer cell lines show similar response to increasing concentration of doxorubicin leading to similarity in overall dynamics **D** cell line-specific estimates of the birth, death, and net growth concentration response curves which are estimated simultaneously in a hierarchical model, borrowing information across cell lines. **E** Estimation of rate specific parameters over all cell lines in each drug independently results in statistics for potency (NG_75_) and efficacy (dNG) that can be used to compare drugs based on their response while accounting for the cell line to cell line variability. Drugs are ranked by the population mean for that statistic and each cell line’s statistic is as well to show the between cell line variability in response which will affect prediction of new cell line responses. **F** Scatterplot of the average NG_75_ and dNG to show the drugs with the greatest response in the top left corner.

We first set out to investigate cell response to doxorubicin (Fig. 4C). We observed that the slopes of the log-cell count over time were visually similar across cell lines (Fig. 4C), justifying the use of a hierarchical model. Previous studies and the Genomics of Drug Sensitivity in Cancer (GDSC) database reported a geometric mean IC_50_ of 0.336 μM for doxorubicin in breast cancer cell lines with a concentration range between 0.004 and 1.02 μM^2^. In contrast, our dataset showed an average 96-hour IC_50_ of 0.018 μM (range: [0.004,0.024] μM). This discrepancy highlights inconsistencies when using IC_50_ values due to differences in experimental conditions and reiterates the need for methods invariant to such variability.

Since we observed continued cell growth even at the highest concentrations tested, we determined two metrics to avoid extrapolation beyond the tested concentration range. First, we defined the NG_75_ as the concentration at which the growth rate is 75% of the control. Second, the dNG is the net difference in growth rate between the control and the largest dose. We found that the average 96-hr IC_25_, the concentration at which 25% growth inhibition is observed (equivalent to 75% of the control) had a mean of 9.4×10^-3^ μM (range: [2.2×10^-3^, 7.1×10^-2^]). The BESTDR-determined average NG*_75_* across cell lines is 7.5×10^-3^ μM (range: [2.5×10^-3^, 1.5×10^-2^]) and the average dNG is 0.020 hr^-1^ (range:[0.008, 0.036], Fig. 4D). These findings demonstrate that the potency values are similar to those from viability assays while efficacy is unaffected by experiment duration. The BESTDR-estimated curves (Fig 4D) indicate that the primary response of breast cancer cell lines to doxorubicin over the concentration range tested is an increase in the death rate, while the birth rate remains nearly constant, suggesting that doxorubicin’s mechanism of action is primarily cytotoxic^41^.

To illustrate how BESTDR can be used to rank drugs for further investigation in high-throughput drug screening experiments, we compared the potency and efficacy of 25 drugs across multiple cell lines. To account for cell line-specific effects, we used the NG_75_ and dNG statistics as determined by BESTDR (Fig. 4E). Drugs with lower NG_75_ values, such as panobinostat, reflect greater potency while drugs with higher dNG values reflect greater efficacy. The range of NG_75_ estimates for the grand mean across cell lines was 5.9 x 10^-l4^ (panibinostat) to 1.9x 10^-5^ (alpelisib). We then investigated the relationship between the two statistics, NG_75_ and dNG, for individual drugs across cell lines (Fig. 4F). The hierarchical model allows calculation of an average response across all cell lines by using the mean of the parameter distributions for each rate to construct concentration-response curves (Fig. 4F). Additionally, we examined mechanism-specific responses across cell lines by defining the metrics B_75_ and dB for birth rates, and D_75_ and dD for death rates which refer to the concentration at which each rate is 75% of the control and the net difference between the control and largest dose, respectively (Fig. S2A,B). For example, volasertib displayed a high dB and low B_75_, indicating a primarily cytostatic effect by inhibiting cell division (Fig. S2A). In contrast, doxorubicin showed a high dD and low D_75_, suggesting a predominantly cytotoxic response through increased cell death (Fig. S2B).

To investigate the generalizability of BESTDR across high-throughput screens, we then analyzed data from two additional studies: one involving PC9-derived non-small cell lung cancer cell lines across 14 drugs (Fig. S2C,D) and another with six small cell lung cancer (SCLC) cell lines across 138 agents (Fig. S2E,F)^33,42–44^. In the PC9 dataset, we observed minimal heterogeneity in drug response among cell lines, likely because they originated from the same parental cell line. The SCLC dataset exhibited greater heterogeneity, emphasizing the utility of using the mean response to account for variability across cell lines. Our analyses demonstrate that drug response data can be effectively summarized using scatterplots combining efficacy and potency metrics. For instance, in the PC9 study, paclitaxel emerged as the most effective drug, while seliciclib was the least effective (Fig. S2D). In the SCLC dataset, most drugs clustered with low dNG and high NG_75_ values, indicating low efficacy and potency but certain drugs, such as SCH-1473759, showed potential for further investigation (Fig. S2F).

These examples illustrate how hierarchical modeling with BESTDR can generate robust estimates by incorporating data from multiple experiments, even when conducted under different conditions. Since the results are based on cell-intrinsic properties independent of experimental variables like time or initial cell count, BESTDR provides consistent and reliable insights for drug ranking and selection in high-throughput drug screening experiments.

### Estimating Transition Rates Between Cell States across Drugs

Since many drugs act specifically on cell state transitions or phases of the cell cycle, we next sought to demonstrate the ability of BESTDR to estimate mechanism-specific drug responses in multi-state systems. We utilized a simplified cell cycle dynamics model that distinguishes cells in the G1 and S/G2/M phases of the cell cycle, using a human DNA helicase B(HDHB) cell cycle reporter^45^. We defined the G1 to S/G2/M transition rate as u_1_(c) and the reverse transition as u_2_ (c), while the death rates of cells in the G1 and S/G2/M phases of the cell cycle are denoted as d_1_(c) and d_2_(c), respectively (Fig. 5A, SI Section 1.5). To validate BESTDR’s ability to accurately estimate parameters in such a system, we simulated data for cell division and phase transitions under single and multiple dose scenarios for estimation of the parameters (SI, Section 1.5) and compared estimates to the ground truth of input parameters. These analyses demonstrate that BESTDR accurately estimates death and transition rates between cell cycle phases (Fig. 5B-E). We found that average cell-count trajectories using the posterior means accurately fit our input data, confirming that BESTDR estimates can be used for prediction (Fig. 5B). In the single-dose scenario, the peaks of the posterior distribution for the estimated rates closely matched the original parameter values (Fig. 5C). Additionally, deterministic trajectories of the posterior mean estimates are fully contained in the synthetic cell count trajectory ranges for each concentration (Fig. 5D). In multiple-dose cases, BESTDR successfully reproduced the original parameters of the logistic function used to describe the concentration-response relationship for each rate (SI, Section 1.4) in that the original parameter values fall within the 95% credible intervals of the estimates (Fig. 5E).

**Figure 5.**
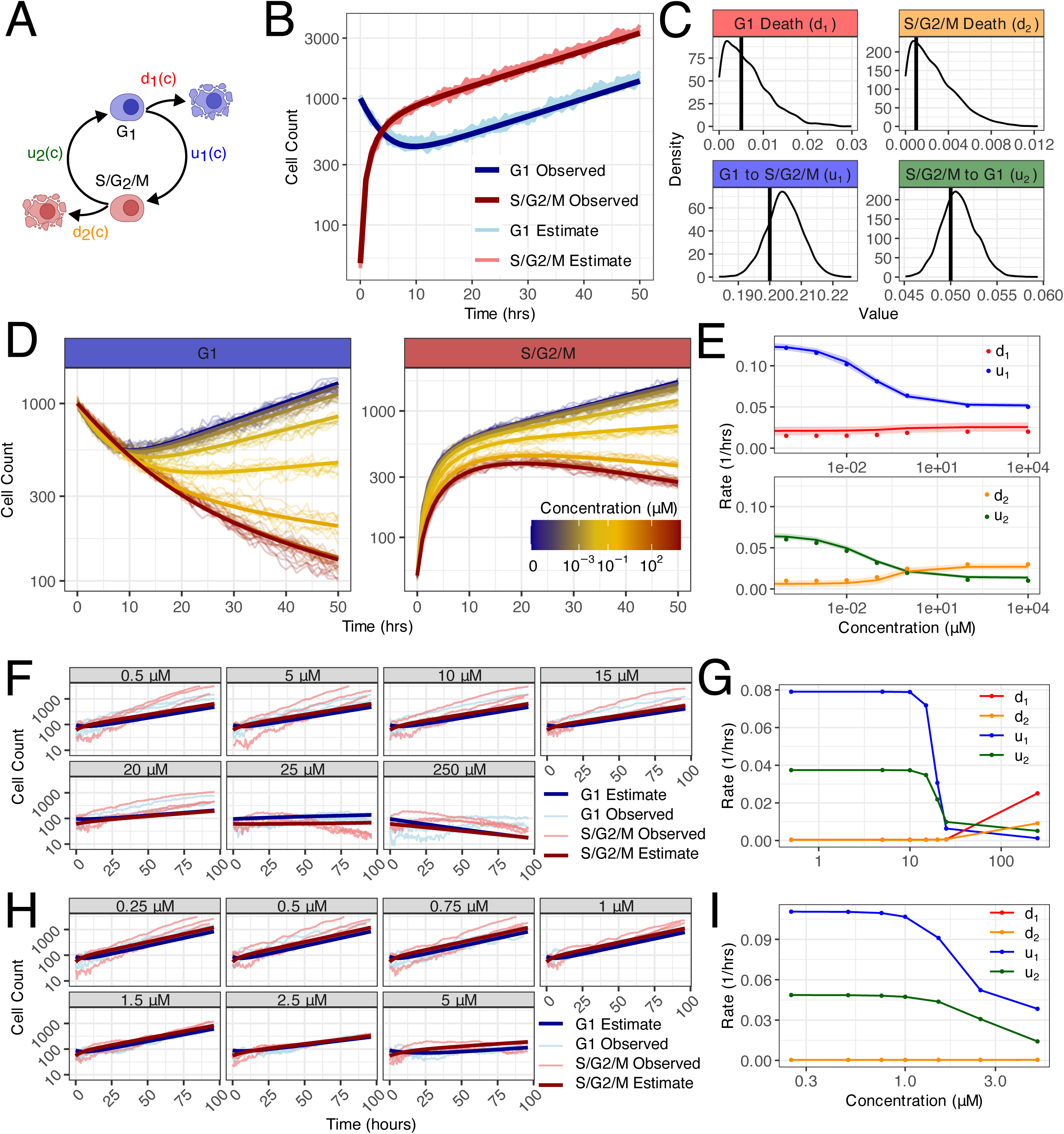
Extending our framework to the cell cycle model allows us to capture the effects of different drugs on transition rates between cell cycle phases. **A** Diagram of the cell cycle model. Here, we consider the G1 phase (blue) and the combined S/G2/M phase (red). With c representing drug concentration, we define the transition rates from G1 to S/G2/M and *vice versa* as u_1_(c) and u_2_ (c), respectively, and the death rates for cells in the G1 and S/G2/M phases as d_1_(c) and d_2_(c), respectively. **B** Synthetic data (light-colored lines) and deterministic estimates (dark-colored lines) obtained by simulating the cell cycle model under a single concentration, using as parameter values the mean of the estimated parameters. **C** Posterior distributions of the estimated parameters. The black solid line indicates the parameter values used to generate the synthetic data. **D** Synthetic data (light-colored lines) and deterministic trajectories (dark-colored lines) obtained by simulating the cell cycle model at multiple concentrations, using as parameter values the mean of the estimated parameters. **E** Estimated parameters for the multiple-dose case using synthetic data using eight different concentration values. Parameter values used to generate the synthetic data (dots) and estimated parameters, represented with their mean (solid line) and 95% credible interval (shade). **F** Observed and estimated trajectories of cell cycle-specific counts under seven concentrations of doxorubicin. **G** Doxorubicin concentration-response curves for each of the cell-cycle rates estimated at each of the observed concentrations. **H** Observed and estimated trajectories of cell cycle-specific counts under seven concentrations of gemcitabine. **I** Gemcitabine concentration-response curves for each of the cell-cycle rates estimated at each of the observed concentrations.

We then applied BESTDR to data from in vitro experiments of cells treated with doxorubicin and gemcitabine at various concentrations counted every 30 minutes for 96 hours^45^. We found that the estimated cell-count trajectories closely matched the average observed dynamics (Fig. 5F,H). Analysis of posterior estimates revealed that higher drug concentrations reduced transition rates u_1_(c) and u_2_(c) for both drugs.

Additionally, higher doxorubicin concentrations increased death rates d_1_(c) and d_2_(c) up to 58-fold and 29-fold, respectively, whereas gemcitabine increased d_1_(c) and d_2_(c) by only 1.11-fold and 1.16-fold (Fig. 5G,I). These results highlight BESTDR’s robustness when modeling complex processes such as cell cycle dynamics under varying drug conditions and its ability to differentiate a drug’s mechanism of action such as inducing apoptosis or inhibiting cell cycle progression.

### Quantifying Reprogramming Dynamics with BESTDR

Finally, we used BESTDR to estimate dynamics of induced cellular reprogramming, where differentiated cells convert into induced pluripotent stem cells (iPSCs) through the constant overexpression of transcription factors Oct4, Klf4, Sox2, and cMyc (OKSM)^14^. Differentiated cells (denoted as D) and iPSCs (denoted as SD) each have their own birth rates (b_D_ (c) and b_SD_(c)) and death rates (d_D_ (c) and d_SD_(c)) as well as an irreversible transition from D to SD cells at rate r(c) (Fig. 6A and SI Section 1.6)^14,46,47^.

**Figure 6.**
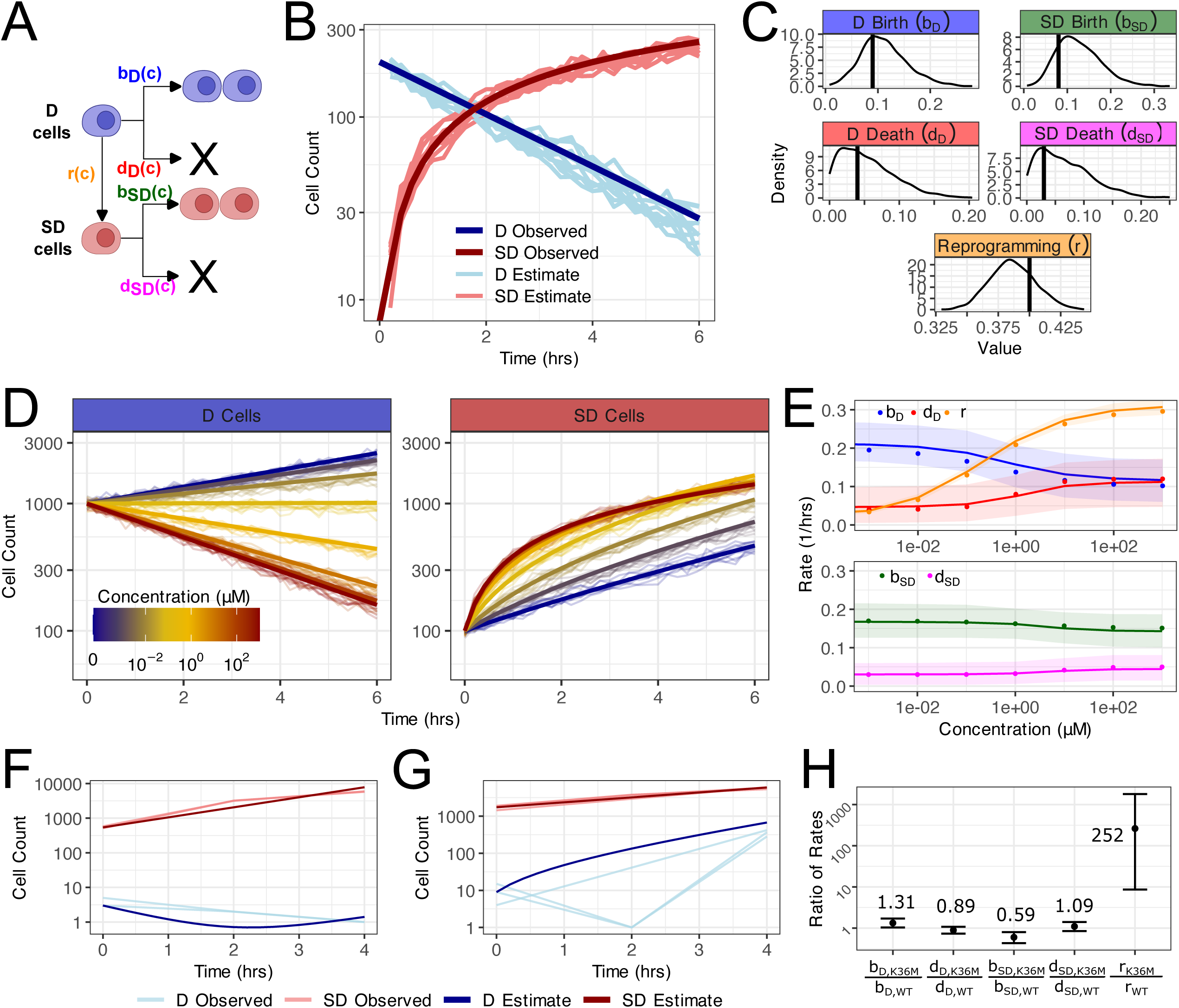
Extending our framework to the reprogramming model allows us to compare the effectiveness of reprogramming approaches of different types of cells. **A** Diagram of the reprogramming model with differentiated cells (D, blue) and iPSC cells (SD, red). With c representing drug concentration, we define birth rates as b_1_(c) and b_2_(c), death rates as d_1_(c) and d_2_(c) and the rate of the reprogramming process as r(c). **B** Synthetic data (dark) for estimation and deterministic trajectories (light) simulated from parameter estimates under the reprogramming model using BESTDR to estimate the specific rates at a single concentration. **C** Posterior distributions of the estimated parameters. The black solid line indicates the parameter values used to generate the synthetic data. **D** Synthetic data (dark) for estimation and deterministic trajectories (light) simulated from parameter estimates under the reprogramming model using BESTDR to estimate the specific rates under a concentration-response curve for each of the five parameters for eight concentrations. **E** True parameter values used to generate the synthetic data (dots) and estimated parameters, represented with their mean (solid line) and 95% credible interval (shade). **F** Observed experimental data (light) and deterministic trajectories (dark) obtained from estimates in wild-type cells with OSKM over four hours. **G** Observed experimental data (light) and deterministic trajectories (dark) obtained from estimates in K36M mutant cells with OSKM over four hours. **H** Estimates of the ratio between the value estimated for the cells with the K36M mutation and the value estimated for the WT cells.

To validate BESTDR’s accuracy when scenarios such as reprogramming dynamics or non-reversible transitions, we simulated data for single-dose and multiple-dose scenarios assuming an increasing transition/reprogramming rate as concentration increases. We used a four-parameter logistic function for each rate for a total of 20 parameters (SI Section 1.6). Unlike standard ODE models (SI Equation 1.5) that can only estimate net growth rates, BESTDR successfully estimated all individual birth, death, and transition rates: the posterior means closely matched the input parameter values and were all contained within the 95% credible intervals for both single-dose (Fig. 6B, C) and multiple-dose scenarios (Fig. 6D,E).

We then applied BESTDR to experimental data of cells with and without the K36M mutation, which inhibits H3K36 methylation, significantly altering chromatin structure and transcriptional regulation and leading to faster reprogramming and a higher proportion of iPSCS relative to wild-type (WT) cells^14^. Cell counts for non-reprogrammed (D) and reprogrammed (SD) cells were quantified via flow cytometry using an Oct4-GFP reporter^14^. We used BESTDR to estimate the birth, death, and reprogramming rates of WT cells (Fig. 6F) and K36M mutant cells (Fig. 6G). The estimated rates were validated by comparing the deterministic cell count trajectories from the mean posterior estimates to the observed data (Fig. 6F,G). We found that the average trajectories were fully contained within the range defined by the observed cell counts, demonstrating alignment between the estimates and observed results (Fig. 6F,G).

Comparing WT and K36M-mutant cells, we found that birth and death rates were similar, with the ratio of the rates for mutated to WT cells ranging from 0.6 (95% CI: [0.43,0.79]) to 1.33 (95% CI: [1.03,1.72], Fig. 6H). However, the K36M mutation greatly increased the reprogramming rate 252-fold (95% CI: [8.36,1806.34], Fig. 6H). Again, trajectories generated from BESTDR estimates closely aligned with the experimental cell count data^14^, demonstrating that approximately 85% of K36M-mutant cells upregulated the pluripotent state reporter between days 4 and 8, compared to 5% of WT cells (Fig. 6H). BESTDR estimates revealed that the K36M mutation primarily increases the reprogramming rate with minimal effects on birth and death rates. These findings demonstrate BESTDR’s utility in quantifying specific effects of the K36M mutation on reprogramming dynamics via OKSM overexpression.

## Discussion

Measuring cell responses to drugs is essential in preclinical studies and drug design to understand mechanisms such as cytotoxic versus cytostatic effects and cell cycle phase-specific actions. As new experimental methods allow observation of finer details and phenotypes in cell culture, new metrics beyond viability are needed to quantitatively assess the impacts of drugs on cell behavior and growth dynamics. Here we introduced BESTDR, a novel Bayesian methodology estimating concentration-response relationships in complex cell growth systems using longitudinal cell count data. BESTDR allows customization of models and concentration-response components, enabling robust estimation across multiple cell lines in hierarchical models suitable for large high-throughput screens. Its flexibility supports modeling complex dynamics, including cell cycling kinetics and state transitions, providing a more comprehensive view of drug responses than traditional metrics. Furthermore, BESTDR’s outputs offer valuable early insights to guide future experiments, such as selecting drugs for combination therapies based on mechanism-specific effects.

Drug dose-response modeling is highly dependent on data quality and can be influenced by errors during data generation. Although BESTDR mitigates some of these errors by incorporating an additional estimation term, high observation error may still lead to slightly inflated rate estimates because observation error cannot be fully separated from the inherent variability in the stochastic processes underlying BESTDR. Advances in computational cell tracking and counting methods, such as artificial intelligence (AI)-based tracking, could help reduce these errors^48–50^. Additionally, Bayesian estimation in BESTDR is sensitive to the choice of priors and sample size, which can affect the accuracy and reliability of the results. When modeling cell cycle dynamics, we observed that the time spent in each cell cycle phase may not follow an exponential distribution, and transient oscillatory behaviors can occur early in experiments until cells reach a stable distribution, as observed in the data^45,51^. Methods based on continuous-time Markov branching processes exist to alleviate these issues by employing the linear-chain trick, thereby allowing BESTDR to model non-exponential times at the cost of increased model complexity^49,52–54^. Finally, high-throughput drug screens sometimes do not capture the full concentration range where drug response may occur, leading to flat concentration response curves and inaccurate potency metrics such as the NG_50_. In these cases, different estimators such as NG_75_ are useful for providing a measure of potency without extrapolating from the model when the NG_50_ is not observed. Note that this problem exists in any concentration-response modeling framework and can be prevented through pilot experiments for dose-finding.

BESTDR provides a versatile and robust framework for dose-response modeling that leverages readily available data to yield insights into mechanisms of drug response. AI and machine learning (ML) have revolutionized data analysis across scientific disciplines including pharmacology and complement more traditional modeling methods by providing robust preprocessing and feature extraction so that models like BESTDR can make robust estimates about more complex systems. Algorithms that automate the extraction of phenotypic features from live-cell imaging, such as morphology-based methods^49^, allow definition of cell states or types that can be used as inputs to BESTDR to model how drugs affect cell morphologies. Further, BESTDR creates an interpretable model maintaining transparency and explainability between AI-derived, “black box” features and cellular responses. We foresee BESTDR being used as part of a hybrid modeling pipeline that combines AI-driven feature extraction from cell morphologies and multi-omics with the mechanistic framework offered by BESTDR to provide deeper insights into cellular response and more robust predictions. By integrating with AI-driven feature extraction and accommodating complex cellular dynamics, BESTDR paves the way for more informed experimental designs and innovative drug development strategies. This holistic approach not only enhances our understanding of cellular responses but also holds promise for accelerating the discovery of effective therapeutic interventions.

## Methods

### Estimation of branching process parameters

The likelihood of a d-type CTMBP at time t started by N individuals at time 0 is approximated by a d-dimensional multivariate normal distribution. This normal likelihood is justified because, in a CTMBP with a sufficiently large initial population, the distribution of cell counts at any given time approximates a normal distribution according to the Central Limit Theorem^23,31^. Given the starting individuals, time, and structure of the branching process, the mean and variance are numerically solved. Due to the Markov property and time-homogeneity assumption, when multiple timepoints exist for the same trajectory, these are split up to act as individual samples, increasing the effective number of samples used to estimate the parameters.

The mean and variance at some time and a given concentration are both solved as functions of the starting number of individuals at the previous time, the estimates for all rates at that concentration, and the length of time between observations. Since the variance is calculated from the stochastic process rather than representing a noise term, the variance in the data serves to help estimate additional parameters that would be unidentifiable using ODE-based methods that do not account for the variance. For example, populations with faster division and death rates exhibit higher variance in cell counts than those with the same net growth rate but slower rates; we leverage this variance to distinguish these rates.

When different drug concentrations are present in the data, a concentration-response function f*_θ_*(c) can be assumed for rate parameter 0 which we assume has parameters *θ_1_*, *θ_2_*,…, *θ*_k_. BESTDR estimates each of these parameters to provide an estimated concentration-response curve for each rate parameter. The likelihood function is updated in this case by writing the mean and variance as functions of drug concentration. We use the 4-parameter logistic function for modeling the concentration-response for each rate, although BESTDR is flexible to any statistical model. For a rate parameter, 0, the 4-parameter logistic function is defined by 0_0_ (the rate at concentration 0), *θ*_inf_ (the rate as the concentration goes to infinity), *θ*_50_ (the log-concentration when the rate is at the midpoint of *θ*_0_ and *θ*_inf_), and *θ*_h_ (the Hill coefficient, or slope at *θ*_50_. We enforce that all rates should be positive by defining d0 as the change between min (*θ*_0_, *θ*_inf_) and max (*θ*_0_, *θ*_inf_). That is if the function is decreasing, then we estimate *θ*_inf_ and d0 and define *θ*_0_ = *θ*_inf_ + d0. Priors are provided for each of the four parameters of the curve from a zero-truncated normal distribution for the *θ*_0_/*θ*_inf_, d0, and 0h and a normal distribution for *θ*_50_. BESTDR uses HMC to estimate posterior distributions for each of the parameters of each concentration-response function. More mathematical details are provided in the Supplement.

### Simulation of in silico experiments

Simulations were performed using the R package *estipop* (v0.0.1)^23^. For each of the simulations we defined parameters representing the rates in single-dose studies or the parameters of the curve in concentration-response studies. When modeling concentration-response curves we generated rates at each tested drug concentration and generated cell count trajectories according to the underlying mechanistic process to use as data for estimation.

### Estimation of Rates

Inference was performed using Stan (v2.34.1)^55,56^ in the R package cmdstanr (v0.7.1)^57^.

### Data Preprocessing and Analysis

Data preprocessing was performed in R (v4.3.2)^58^ using the *tidyverse* (v2.0.0)^59^ for data manipulation and plotting.

### Cell Culture Experiments

HCT-116 p53-VKI cells (Lahav lab, Harvard Medical School) were seeded in 96 well plates and imaged using an Incucyte S3 imaging platform. Cisplatin was added at increasing concentrations (maximum concentration of 25μM with a 2-fold increase for 7 dose points) 24hrs following cell seeding. Multiwell plates were imaged every 4hrs over a period of 72hrs following addition of cisplatin. Dead cells were monitored using staining for Ethidium Bromide. Imaging was performed using a 10X objective to identify the total number of cells using the green fluorescent protein channel (GFP, 300ms acquisition time) and dead cells using the red fluorescent protein channel (RFP, 400ms acquisition time). The top-hat segmentation algorithm of the Incucyte S3 software platform was used to detect objects including GFP+ (total cells), and GFP+RFP+ (dead cells). We set the parameters for the GFP+ objects at a radius 30μm, an intensity threshold of 0.1 and filtered for objects smaller than 100μm^2^, while the parameters for the RFP+ were set at a radius of 15μm, an intensity threshold of 0.05 and filtered for objects smaller than 30μm^2^.

## Acknowledgements

This research is supported by National Institutes of Health (NIH)/National Institute of General Medical Sciences grants R01GM144962 to T.O.M. and R35GM150815 to I.K.Z. We gratefully acknowledge support of the Ludwig Center at Harvard and the Dana-Farber Cancer Institute’s Center for Cancer Evolution.

## Contributions

T.O.M and F.M. conceived the study. T.O.M. and J.P.R. developed the method and wrote the code. T.O.M. and S. B. performed analysis on all simulated and experimental data. T.O.M., S.B., and F.M. wrote the manuscript. I.K.Z. performed cell culture experiments and performed analysis on experimental data. All authors reviewed and approved the manuscript.

## Disclosures

F.M. is a co-founder of and has equity in Harbinger Health, has equity in Zephyr AI, and serves as a consultant for both companies. She is also on the board of directors of Recursion Pharmaceuticals. F.M. declares that none of these relationships are directly or indirectly related to the content of this manuscript.

---

**Figure S1.**
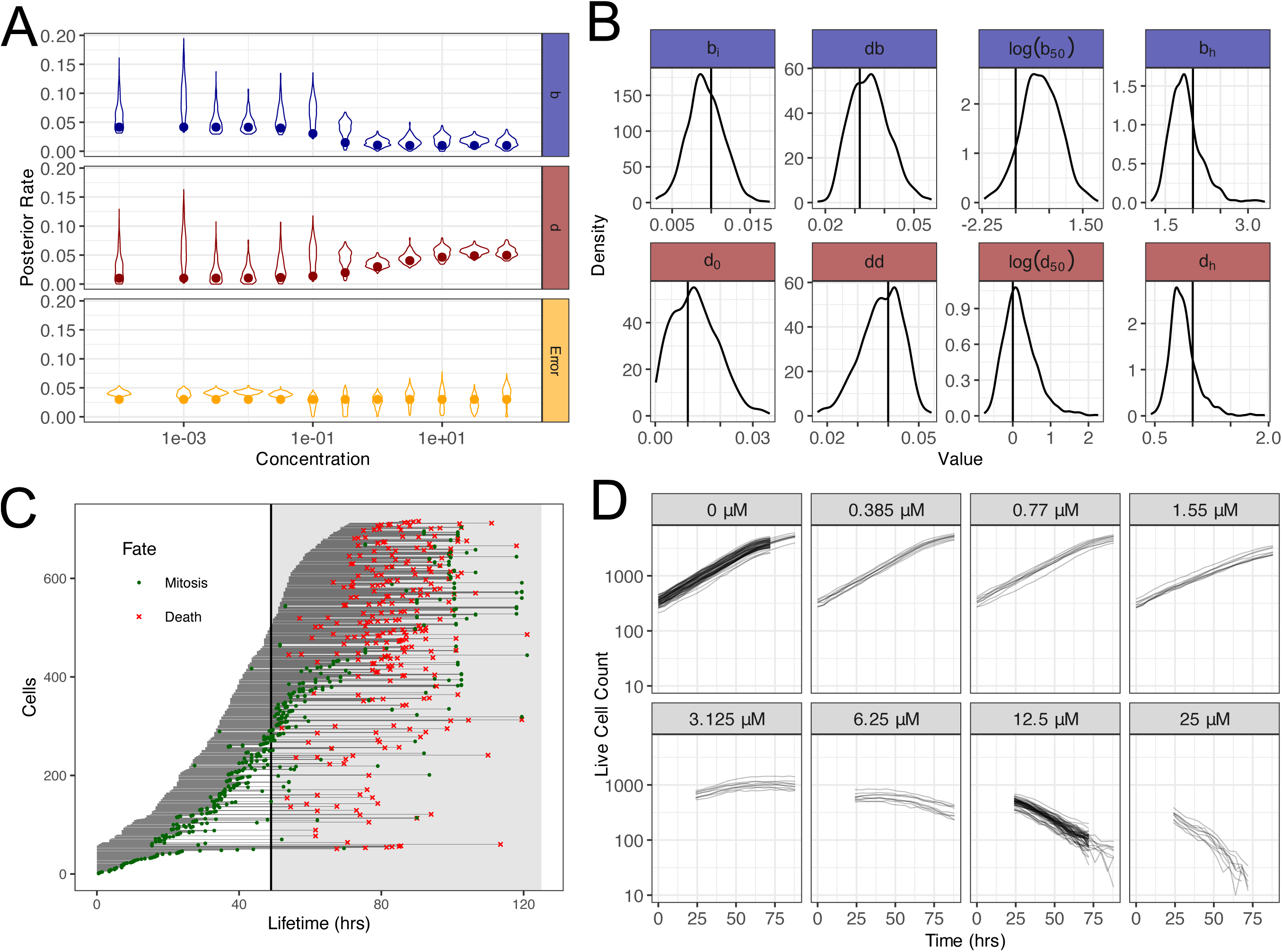
BESTDR estimation of rates in silico and in vitro live cell counting experiments. **A** Individual posterior estimates from simulated data for the birth and death rate in the violin plot along with the true value when each concentration is estimated separately without a concentration-response curve. **B** Posterior estimates for the parameters of the concentration response curves along with their true values from the simulated data in a concentration-response model. **C** Individual cell fates from the cell fate tracking where cells either split (green) or die (red). The length in the x-axis determines the time and the cell fate determines which distribution it counts toward when building the validation time-distribution to compare to the estimates from BESTDR. **D** Cell count data over multiple replicates at various levels of cisplatin are used to build a concentration-response for the birth and death rates of HCT-116 p53-VKI.

**Figure S2.**
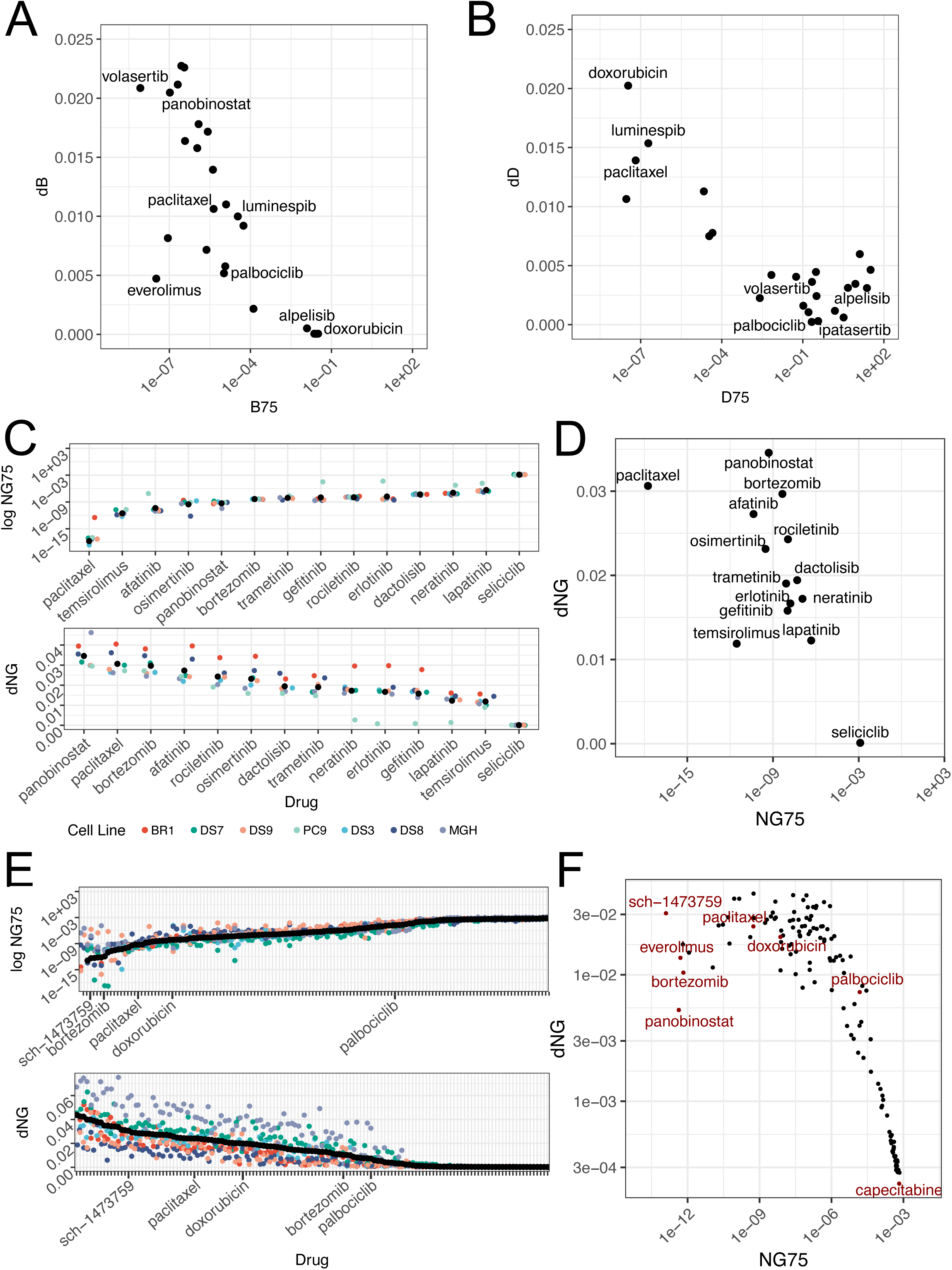
Hierarchical modeling allows simultaneous comparison of many drugs in screens while accounting for variability across cell lines. **A** Scatterplots of the birth (left) rate concentration-response statistics for potency (B_75_, x-axis) and efficacy (dB, y-axis) provide insight into the cytostatic mechanistic effects of each drug. **B** Scatterplots of the death (left) rate concentration-response statistics for potency (D_75_, x-axis) and efficacy (dD, y-axis) provide insight into the cytotoxic mechanistic effects of each drug. **C** Hierarchical modeling of 7 PC9 sub-cell lines across 14 agents with the NG_75_ and dNG ranked. **D** Scatterplot of the NG_75_ and dNG to show both drug potency and efficacy in 7 PC9 sub-cell lines. **E** Hierarchical modeling of 6 small-cell lung carcinoma cell lines across 138 agents with the NG_75_ and dNG ranked (left) and as a scatterplot (right). **F** Scatterplot of the NG_75_ and dNG to show both drug potency and efficacy in 6 small-cell lung carcinoma cell lines.

