## Supplemental Information for "BESTDR: Bayesian quantification of mechanism-specific drug response in cell culture"

### Supplementary Information to *Quantification of mechanism-specific drug response in cell culture*

December 23, 2024

### 1 Extended Methods

#### 1.1 Single-Concentration Branching Process Estimation

In an experiment containing $d$ different cell types that interact, we model the number of cells of each type as a $d$-type continuous-time Markov branching process (CTMBP), counted by the vector $\mathbf{Z}(t;\mathbf{N}) = (Z_1(t;\mathbf{N}), Z_2(t;\mathbf{N}), \ldots, Z_d(t;\mathbf{N}))$ given $\mathbf{N} = (N_1, N_2, \ldots, N_d)$ cells of each type at time 0. Let $u_{i\mathbf{j}} = (Z_1(t;\mathbf{N}), Z_2(t;\mathbf{N}), \ldots, Z_d(t;\mathbf{N}))$ be the rate parameter associated with the type $i$ parent cell begetting $\mathbf{j} = (j_1, j_2, \ldots, j_d)$ children. As an example, the simple birth-death process counts cells of a single type, $\mathbf{Z}(t;N)$, and the birth and death rates are $b \equiv u_{1(2)}$ and $d \equiv u_{1(0)}$.

We have previously shown that given a sufficiently large number of ancestors, the branching process, $\mathbf{Z}(t;\mathbf{N})$, is approximately normally distributed where the mean vector and covariance matrix are described by first-order ODEs that can be numerically solved given the parameters for the process [1]. For brevity, we write the mean vector and covariance function of the process as $\mathbf{M}(\mathbf{u}) \equiv \mathbf{M}(t;\mathbf{N},\mathbf{u})$ and $\mathbf{\Sigma}(\mathbf{u}) \equiv \mathbf{\Sigma}(t;\mathbf{N},\mathbf{u})$. Thus we approximate the distribution of $\mathbf{Z}$ as

$$\mathbf{Z}(t;\mathbf{N}) \sim N(\mathbf{M}(\mathbf{u}), \mathbf{\Sigma}(\mathbf{u})).$$

Assuming $m$ realizations of the branching process, $\mathbf{z}(t) \equiv [\mathbf{z}^{(i)}(t^{(i)};\mathbf{N}^{(i)})]_{i=1,\ldots,m}$, where $t^{(i)}$ represents the time and $\mathbf{N}^{(i)}$ represents the ancestor vector of the $i^{\text{th}}$ realization, we further define $\mathbf{M}^{(i)}(\mathbf{u}) \equiv \mathbf{M}(t^{(i)};\mathbf{N}^{(i)},\mathbf{u})$ and $\mathbf{\Sigma}^{(i)}(\mathbf{u}) \equiv \mathbf{\Sigma}(t^{(i)};\mathbf{N}^{(i)},\mathbf{u})$ as the mean and variance functions for the $i^{\text{th}}$ realization given the parameters, $\mathbf{u}$. We note that $t$ and $\mathbf{N}$ do not have to be identical for each realization and for nonidentical values we can resolve the ODEs for the moments.

the log-likelihood function for the rate parameters $\mathbf{u}$ is then the normal log likelihood function,

$$\log L(\mathbf{z}(t);\mathbf{u}) \propto -\frac{m}{2}\log|\mathbf{\Sigma}^{(i)}(\mathbf{u})| - \frac{1}{2}\sum_{i=1}^{m}[\mathbf{z}^{(i)}(t) - \mathbf{M}^{(i)}(\mathbf{u})]^T\Sigma^{(i)}(\mathbf{u})^{-1}[\mathbf{z}^{(i)}(t) - \mathbf{M}^{(i)}(\mathbf{u})].$$

Given the parameter vector of rates, $\mathbf{u}$, we seek the posterior distribution, $p(\mathbf{u}|\mathbf{Z}) \propto \pi(\mathbf{u})L(\mathbf{z}(t);\mathbf{u})$ where $\pi(\mathbf{u})$ represents the prior distribution for the rate parameters. We use Markov chain Monte Carlo methods to sample from the posterior density $p(\mathbf{u}|\mathbf{Z})$ using Hamiltonian Monte Carlo with the Stan software package ([2, 3, 4]).

#### 1.2 Concentration-Response Estimation

We further extend the model by considering the situation where the rate parameters are instead a function of additional data such as drug concentration, $c$, and parameters, $\boldsymbol{\theta}$, that define a concentration-response curve, or $\mathbf{u}(c;\boldsymbol{\theta}) = (u_1(c;\boldsymbol{\theta}),\ldots,u_p(c;\boldsymbol{\theta}))$.

As an example, let $u_i$ be rate parameter that is sigmoid function of the log-concentration following the 4-parameter logistic function,

$$u_i(c;\theta_0,\theta_{inf},\theta_{50},\theta_H) = \theta_{inf} + \frac{\theta_0 - \theta_{inf}}{1 + \theta_H(log(c) + \theta_{50})}$$

where

$$\begin{aligned}
\theta_0 &= \text{response at 0 concentration} \\
\theta_{inf} &= \text{response at infinite concentration} \\
\theta_{50} &= \text{log concentration at the midpoint of } \theta_0 \text{ and } \theta_{inf} \\
\theta_H &= \text{Hill coefficient.}
\end{aligned}$$

A realization of the process comprises the data $\left(\mathbf{z}^{(i)}(t^{(i)}), t^{(i)}, \mathbf{N}^{(i)}, c^{(i)}\right)$. The likelihood $L(\mathbf{z}(\mathbf{t}); \boldsymbol{\theta})$ becomes a function of the parameter vector, $\boldsymbol{\theta}$, and the moment functions are functions of the rates, $M^{(i)}(\mathbf{u}(c; \boldsymbol{\theta}))$ and $\Sigma^{(i)}(\mathbf{u}(c; \boldsymbol{\theta}))$. The same MCMC method is then used to estimate the posterior for the parameters $\boldsymbol{\theta}$.

The underlying functional form of the concentration-response parameter of the rate can be over multiple drugs and describe the interaction between drugs at various concentrations, $\mathbf{u}(c_1, c_2)$, or even have a nonparameteric form using Gaussian processes.

We further extend this method to hierarchical models where $\boldsymbol{\theta}_j$ is the parameter set for a process of cells coming from group $j$ containing the elements $\boldsymbol{\theta}_j = (\theta_{j1}, \ldots, \theta_{jl_j})$ where for any $j$, $\theta_{jk}$ has has a prior distribution $\pi(\boldsymbol{\gamma_j})$ and $\boldsymbol{\gamma_j}$ represents the hyperparameters that describe the distribution of parameters over all groups. We show such a model is used to provide estimates accounting for data from multiple cell lines to more accurately compare drug responses.

##### 1.3 Observation Error

We model the observation error of the process as a multiplicative factor of the mean at some time, $t$. That is, assuming a random variable, $\epsilon$, representing the error term, we modify the observed cell count distribution's variance to include an additional term that is proportional to the mean and the parameter is constant,

$$\mathbf{Z}(t; \mathbf{N}) \sim N(\mathbf{M}(\mathbf{u}), \boldsymbol{\Sigma}(\mathbf{u}) + \epsilon \mathbf{M}(\mathbf{u})).$$

The value, $\epsilon$, can represent the percent of error between the observed cell count and the total cell count possible due to miscounting of the microscopy software. This additional error adds another degree of freedom and can affect the estimate due to affecting the variance. We have observed for low values of the error though it does not affect the estimation. We expect error rates to be near zero, or a few percentage point, but note that error would be system to all concentration-response modeling.

#### 1.4 Birth-Death Process Concentration-Response Model

Given trajectories of cell growth over time that count the total number of living cells, we with to estimate the birth and death rates as functions of the concentration of drug. For any trajectory counting the number of cells at times $0, t_1, t_2, \ldots, t_m$, we assume the process is time-homogeneous and Markov so that we can break the trajectory into intervals of size $\Delta t_i = t_i - t_{i-1}$. Thus

$$\begin{aligned} Z(t_m|0,1,\ldots,t_{m-1}) &\stackrel{D}{=} Z(t_m|t_{m-1})Z(t_{m-1}|t_{m-2})\ldots Z(t_1|0) \\ &\stackrel{D}{=} Z(\Delta t_m|0)Z(\Delta t_{m-1}|0)\ldots Z(t_1|0). \end{aligned}$$

We treat each interval independently as data using the cell count at the end of the interval as $z$, the cell count at the beginning of the interval as $N$, $\Delta t$ as the time interval, and the log of the concentration as $c$.

For any cell we assume present in log-concentration $c$ of drug, it will either split into two daughter cells after an exponentially distributed length of time with parameter $b(c; b_{inf}, \Delta b, b_{50}, b_H)$ which we assume decreases monotonically with concentration or die after an exponentially distributed length of time with parameter $d(c; d_0, \Delta d, d_{50}, d_H)$ which we assume increases monotonically with increasing concentration. The functional form of equal equation is a 4-parameter logistic function transformed so that we can constrain the rates to be greater than zero. That is

$$b(c; b_{inf}, \Delta b, b_{50}, b_H) = b_{inf} + \frac{\Delta b}{1 + \exp(b_H * (c - b_{50})} \tag{1.1}$$

and

$$d(c; d_0, \Delta d, d_{50}, d_H) = d_0 + \Delta d - \frac{\Delta d}{1 + \exp(d_H * (c - d_{50})} \tag{1.2}$$

where

$$\begin{aligned}
b_{inf} &\equiv \text{birth rate at infinite dose on } (0, \infty) \\
d_0 &\equiv \text{death rate at control on } (0, \infty) \\
\Delta b / \Delta d &\equiv \text{total change in the birth/death rate on } (0, \infty) \\
b_{50} / d_{50} &\equiv \text{the log-concentration at the midpoint of the birth/death rate on } (-\infty, \infty) \\
b_H / d_H &\equiv \text{Hill coefficient of the birth/death rate on } (0, \infty).
\end{aligned}$$

#### 1.5 Cell cycle model: further derivations

The cell cycle model considered in this paper was first proposed in [5]. In this model we lump together phases S, G2, and M in a single phase, called the S/G2/M phase, and we consider only two phases: the G1 phase and the S/G2/M phase. The model can be represented by the circuit diagram shown in Figure 5A.

Defining the count of cells in G1 phase as $x_1$ and the count of cells in S/G2/M phase as $x_2$, let us now introduce the corresponding ordinary differential equation (ODE) model in terms of $x_1$ and $x_2$. It is possible to do this by assuming the total cell count large enough to consider $x_1$ and $x_2$ real numbers. Now, let $c$ represent the log concentration of the drug, let $u_1(c)$ represent the transition rate from G1 phase to S/G2/M phase, let $u_2(c)$ represent the transition rate from S/G2/M phase to G1 phase, let $d_1(c)$ represent the death rate of cells in G1 phase, and let $d_2(c)$ represent the death rate of cells in S/G2/M phase. The ODEs can then be written as

$$\begin{aligned}
\dot{x}_1 &= -u_1 x_1 - d_1 x_1 + 2 u_2 x_2 \\
\dot{x}_2 &= u_1 x_1 - u_2 x_2 - d_2 x_2,
\end{aligned} \tag{1.3}$$

in which, to simplify the notation, we did not explicitly show the dependency of each rate on the concentration $c$.

In the multiple-dose case, we modeled the parameters as 4-parameter logistic functions of the drug concentration $c$. More precisely, we have

$$
\begin{aligned}
u_1(c; u_{1,inf}, \Delta u_1, u_{1,50}, u_{1,H}) &= u_{1,inf} + \frac{\Delta u_1}{1 + \exp(u_{1,H} * (c - u_{1,50})}, \\
u_2(c; u_{2,inf}, \Delta u_2, u_{2,50}, u_{2,H}) &= u_{2,inf} + \frac{\Delta u_2}{1 + \exp(u_{2,H} * (c - u_{2,50})}, \\
d_1(c; d_{1,0}, \Delta d_1, d_{1,50}, d_{1,H}) &= d_{1,0} + \Delta d_1 - \frac{\Delta d_1}{1 + \exp(d_{1,H} * (c - d_{1,50})}, \\
d_2(c; d_{2,0}, \Delta d_2, d_{2,50}, d_{2,H}) &= d_{2,0} + \Delta d_2 - \frac{\Delta d_2}{1 + \exp(d_{2,H} * (c - d_{2,50})},
\end{aligned}
\tag{1.4}
$$

where all the parameters are defined as done in (1.1) and (1.2).

In the first part of the study, we generated synthetic data and used them as inputs for estimation. To ensure biological consistency, we selected parameters that reflect expected cell cycle progression dynamics over time. In the two-phase model, the transition rates between the G1 and S/G2/M phases capture the speed of cell cycle progression, and they typically range from 0.1 to 0.01 per hour, depending on the cell type and environmental conditions [6, 7].

In rapidly proliferating cells, such as cancer cells and stem cells, death rates are generally lower than transition rates. This is because these cells prioritize division over apoptosis, which drives population growth. For instance, studies have demonstrated that division rates are significantly higher than apoptosis rates, supporting tumor expansion. The assumption of lower death rates than transition rates allows the model to realistically capture the behavior of proliferating cell populations [8].

##### 1.6 Switch model: further derivations

The switch model used in this paper to study the cellular reprogramming process includes two types of cells, i.e., type 1 (representing differentiated cells, such as fibroblasts) and type 2 (representing iPSCs), and, as previously assumed in other works ([9, 10, 11]), only the transition from type 1 to type 2, but not *viceversa*. The model can be represented by the circuit diagram shown in Figure 6A.

Defining the count of type 1 cells as $x_1$ and the count of type 2 cells as $x_2$, let us now introduce the corresponding ordinary differential equation (ODE) model in terms of $x_1$ and $x_2$. It is possible to do this by assuming the total cell count large enough to consider $x_1$ and $x_2$ real numbers. Now, let $c$ represent the log concentration of the drug, let $b_1(c)$ represent the birth rate of type 1 cells, let $b_2(c)$ represent the birth rate of type 2 cells, let $d_1(c)$ represent the death rate of type 1 cells, let $d_2(c)$ represent the death rate of type 2 cells, and $r(c)$ represent the transition rate from type 1 to type 2. The ODEs can then be written as

$$\begin{aligned} \dot{x}_1 &= b_1 x_1 - d_1 x_1 - r x_1 \\ \dot{x}_2 &= b_2 x_2 - d_2 x_2 + r x_1, \end{aligned} \tag{1.5}$$

in which, to simplify the notation, we did not explicitly show the dependency of each rate on the concentration $c$.

In the multiple-dose case, we modeled the parameters as 4-parameter logistic functions of the drug concentration $c$. More precisely, we have

$$\begin{aligned} b_1(c; b_{1,inf}, \Delta b_1, b_{1,50}, b_{1,H}) &= b_{1,inf} + \frac{\Delta b_1}{1 + \exp(b_{1,H} * (c - b_{1,50})}, \\ b_2(c; b_{2,inf}, \Delta b_2, b_{2,50}, b_{2,H}) &= b_{2,inf} + \frac{\Delta b_2}{1 + \exp(b_{2,H} * (c - b_{2,50})}, \\ d_1(c; d_{1,0}, \Delta d_1, d_{1,50}, d_{1,H}) &= d_{1,0} + \Delta d_1 - \frac{\Delta d_1}{1 + \exp(d_{1,H} * (c - d_{1,50})}, \\ d_2(c; d_{2,0}, \Delta d_2, d_{2,50}, d_{2,H}) &= d_{2,0} + \Delta d_2 - \frac{\Delta d_2}{1 + \exp(d_{2,H} * (c - d_{2,50})}, \\ r(c; d_0, \Delta r, r_{50}, r_H) &= r_0 + \Delta r - \frac{\Delta r}{1 + \exp(r_H * (c - r_{50})}, \end{aligned} \tag{1.6}$$

where all the parameters are defined as done in (1.1) and (1.2).

#### 2 Combination Models

##### 2.1 Modeling mitosis and death rates as a function of multiple drugs enables us to estimate drug-specific effects for optimal combinations

We considered a model where we treat a cells with a combination of two drugs under the assumption that the birth and death rates are each functions of both drugs (SI Figure 1A). A previous study treated four breast cancer cell lines with a combination of 20 concentrations of paclitaxel over a range of 0 to 4uM with 14 concentrations of carboplatin ranging from 0 to 0.32uM [12]. We estimated the mean normalized viability of the MDA-MB468 cell line growth for each combination relative to the control and observe that the viability over each drug suggests the Bliss independence model serves as a good model for CR (Figure 1B). The Bliss model assumes that the product of the marginal CR curves gives the combination effect and there is no additional synergy between drugs. The predicted viability surface is created from the marginal CR estimates for viability for each drug that was estimated independently, and the predicted viability is shown in SI Figure 1C along with the IC50 (red curve). The similarity between the shape of the surfaces suggests the Bliss model is appropriate to model response.

##### 2.2 Birth and death estimates of paclitaxel and carboplatin in combination

Time course measurements of cell counts that were recorded over about 115 hours were used to estimate the birth and death CR surfaces. We found that in each drug in isolation, the birth rate tended to drop and reach its lower asymptote before the death rate increased, leading to biphasic growth curves. However, we note that the low measured concentrations leads to a large variability in the death rates at higher concentrations of each drug, so the death rates are likely strongly influenced by the prior distributions and drug response is likely driven by decreases in birth rate, or cytostatic effects (SI Figure 1E). The mean posterior b50 (concentration where the birth rate is half that of the control) is 2e-3 uM for paclitaxel and 22uM for carboplatin which appear to line up with the point in the time course data where the cell trajectories experience a sharp decrease.

Using the posterior mean growth rate surface in SI Figure 1F, we show that the IC50 isobole suggest lower concentrations of paclitaxel and similar concentrations of carboplatin required to get a 50% response as our NG50 (yellow curve, SI Figure 1F). The NG0 isobole is also illustrated in green representing the concentration of each drug required to achieve no net cell growth. Based on the posterior predicted surface we observe that carboplatin does not necessarily affect the growth rate of MDAMB468 cells that already have a sufficiently high concentration of paclitaxel. We further estimated the parameters in a mixed effects model using all four cell lines and found a similar result, where paclitaxel had a similar NG50 and NG0 across all four cell lines while carboplatin was much more variable (SI Figure 1F). This is likely due to paclitaxel being the primary driver of response and the marginal curve for carboplatin being flatter.

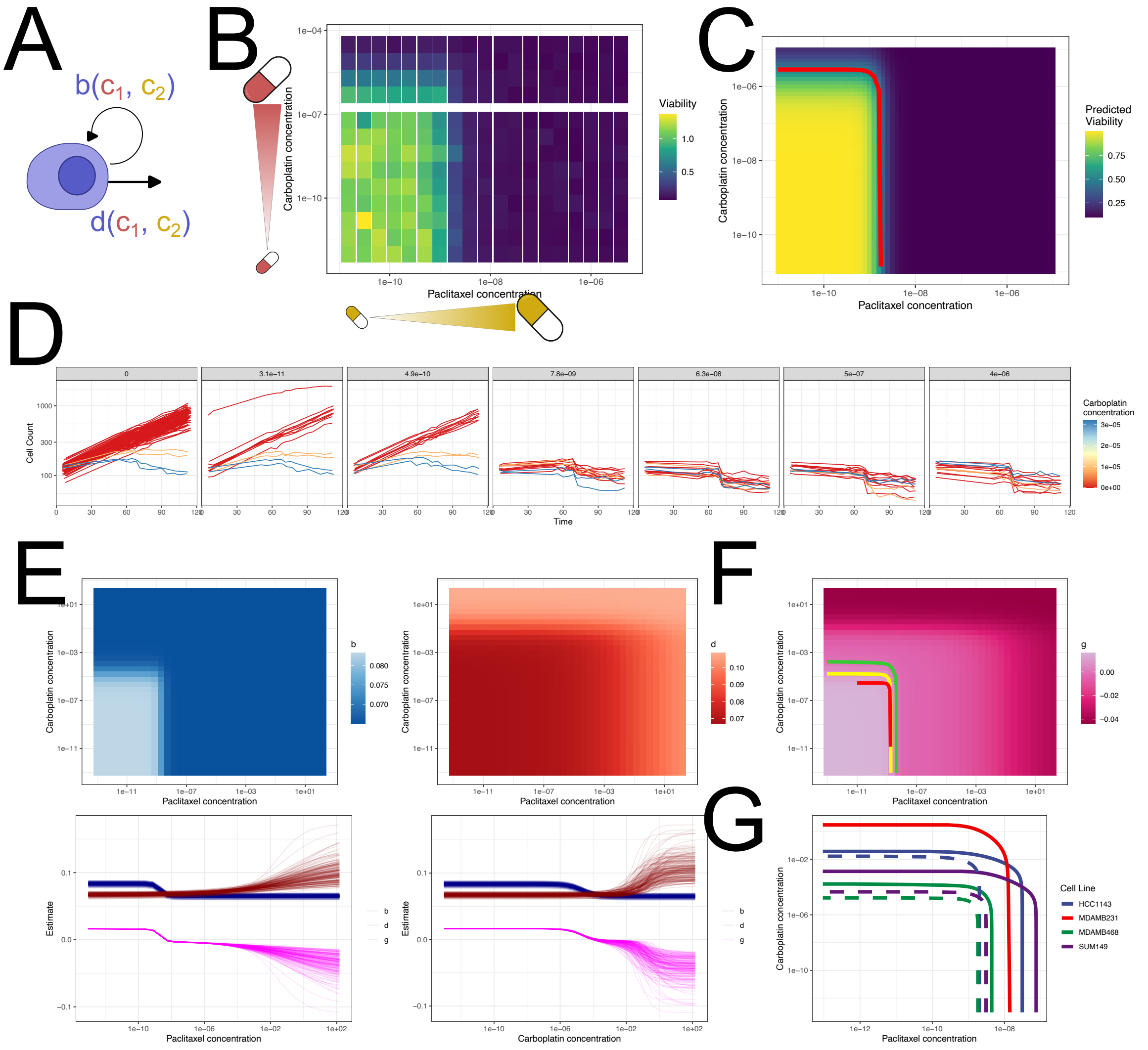

SI Figure 1: **A.)** A combination model of drug response can treat the birth and death rate as functions of more than one drug's concentration. **B.)** Viability of MDA-MB468 cell line in a combination of paclitaxel and carboplatin using cell count relative to the control. **C.)** Predicted relative viability under the assumption of multiplicative effects with the corresponding IC50 curve (red). **D.)** A sample of cell growth data in paclitaxel and carboplatin combinations using live-cell imaging. **E.)** Estimated birth rate (blue) and death rate (red) surfaces using a branching process model shows different concentrations of each drug where the rates change. Across both drugs, the birth rate slows down at lower concentrations before the death rate increases. Realizations from the posterior distribution for the marginal values where one of the drugs is at 0 uM are shown on the bottom. **F.)** The net growth rate surface shows a biphasic change in growth in each direction and G50 (yellow), G0(green) and previous IC50 (red) are shown. **G.)** A hierarchical model of concentration-response over 4 cell lines shrinks the estimates for the cell line specific G50 (dashed) and G0 (solid).

### 3 Analyzing Sample Size

#### 3.1 Sample size effect for single type processes

We create a single-type birth death process to show the effect of the reduction of data on BESTDR's estimation accuracy. We consider a full birth-death process where we simulate branching process growth over 12 concentrations in 8 replicates that we might see when using a 96 well plate. The birth and death rate functions are described by 4-parameter logistic curves that are illustrated in SI Figure 2A. The true cell growth trajectories used as data are shown in SI Figure 2B, and show the growth decrease with increasing concentration and even significant death in the higher concentrations. Like previous experiments, we run the simulation for 72 hours taking cell counts every 4 hours to replicate an live-cell imaging platform. We consider the following scenarios where we reduce the data: (a.) the full data, (b.) 4 of the replicates across all time points, (c.) using only 3 time points (24, 48, 72 hr) but all replicates, and (d.) using only 3 time points and 3 replicates. We also included an error term to make the simulation more realistic, where we added Gaussian noise at a level proportional to the average size of the population at each time point which would represent the percent error in observation from the true cell count.

We use relatively uninformed priors and keep them the same across all scenarios (SI Figure 2C, red dotted line). These priors, while small, have a large enough variance and should cover a large range for possible cell splitting and death rates, so we should not see unreasonably short doubling times as estimates. The true value for each of the parameters is given in vertical black lines and the posterior density is given in solid blue in SI Figure 2C. The full simulation has a peak nearly at the actual value in each of the parameters with the exception of the error term. The dose-response curve also reflects this accuracy and the 95% intervals are small beyond the left asymptote (SI Figure 2D, left). As the data sample size is reduced, the standard error becomes slightly larger, but BESTDR still manages to estimate the curves' parameters. The final scenario shows bimodality though with one of the two modes near the true value. This bimodal effect was previously observed in the 4-parameter logistic dose-response curves for the birth-death process where the $b_{50}/d_{50}$ and $bh/dh$ show the two modes. Because the modes are near eaach other, it does not have a great impact on the predicted dose-response curve, but does increase the error for it.

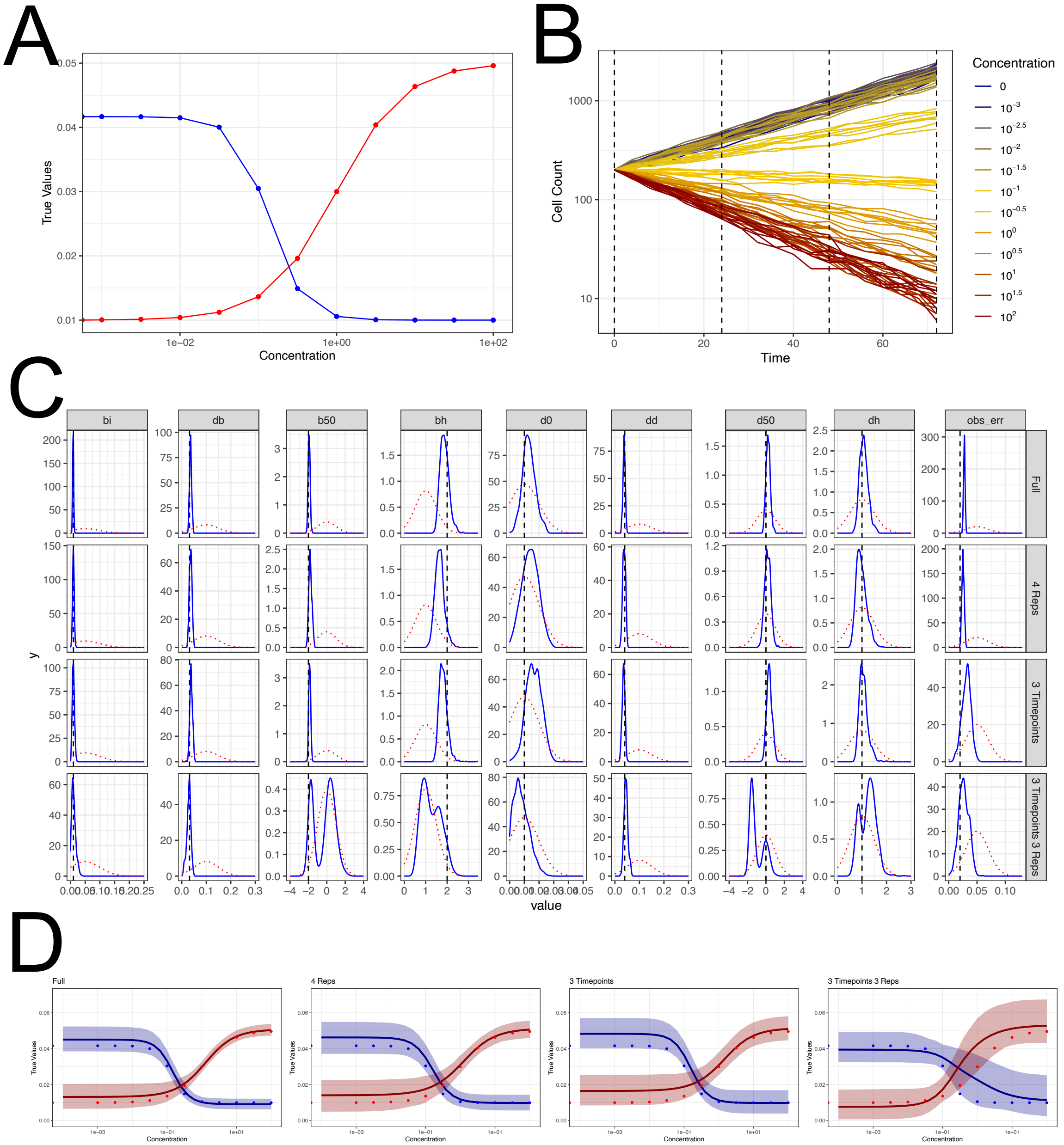

SI Figure 2: **A.)** A concentration-response curve with points indicating concentrations used to simulate data. **B.)** Simulations from each concentration where we tracked cell counts every four hours for 72 hours across 10 replicates. **C.)** Priors (red) and posterior (blue) estimates for each parameter (columns) across the 4 different scenarios (rows). **D.)** Concentration-response curves and credible bands for each of the 4 scenarios using different amounts of data along with the true values as points.

##### 3.2 More accurate priors and larger sample sizes improve parameter estimation in a cell cycle model

We performed a similar test with simulated data for the 2-parameter cell cycle model under a single concentration. We set priors with means equal to the true value or far from the true value as well as having small and large standard deviations to show the effect of the priors. Additionally we changed the number of replicates for sampling. For each case, as a top panel we report the synthetic data (light-colored lines) and deterministic trajectories (dark-colored lines) obtained by simulating our cell cycle model (SI Figure 3), using as parameter values the mean of the estimated parameters. As a bottom panel, we have the posterior distributions of the estimated parameters. The black solid line indicates the parameter values used to generate the synthetic data. We see that badly informed priors (right column) typically lead to estimates far from the true values, but uninformed priors that have a high standard deviation and a mean near the true values perform well. As we increase the data we have posterior estimates with smaller variance.

#### 4 Guide to Software

We included code in the form of an R package that allows users to download and implement BESTDR on their own data (`https://github.com/olliemcdonald/bestdr`). Within the code we have included Stan model files as well as R code for simulating concentration-response data for specific models. Additionally we created an R package that allows the user to define a mechanistic model and the concentration-response curves for the rates which will convert the model into usable Stan code for compilation and estimation.

##### 4.1 Model Library

The model library contains specific models in single dose or with concentration-response relationships between a drug and the rates. These files are under the `/inst/` directory. Users can access these files in R with

```
system.file("model_library/1type-livecell/logistic/birthdeath.stan",
```

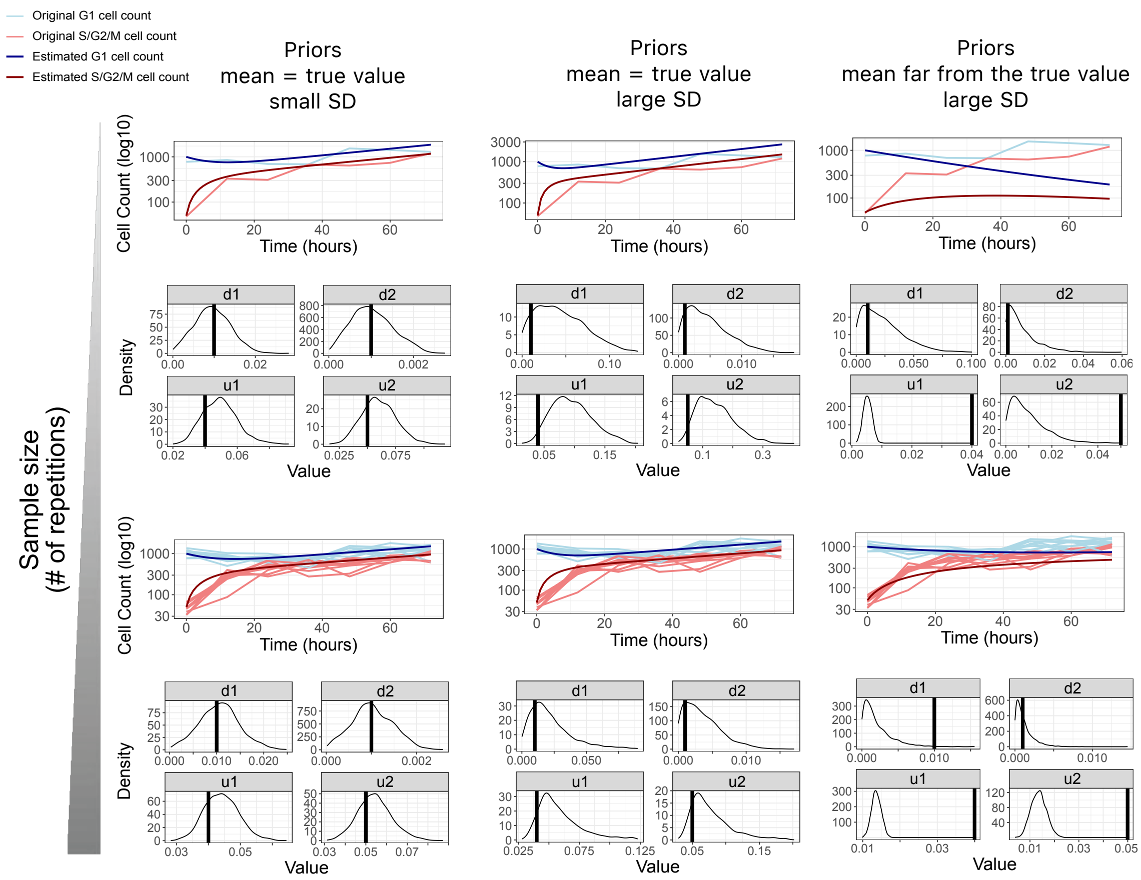

SI Figure 3: Estimated parameters for the cell-cycle model (Figure 5A) - single-dose case, using synthetic data as input and considering different priors and different numbers of repetitions. For each case, as a top panel we report the synthetic data (light-colored lines) and deterministic trajectories (dark-colored lines) obtained by simulating our cell cycle model (SI - Eqs. (1.3)), using as parameter values the mean of the estimated parameters. As a bottom panel, we have the posterior distributions of the estimated parameters. The black solid line indicates the parameter values used to generate the synthetic data.

```
    package = "bestdr")
```

We also included some example scripts that simulate data and implement the estimation to give users a familiarity with the code. The R package additionally includes a vignette for simulating and running code for a simple example that follows an example script.

##### 4.1.1 Models in the Model Library

1. 1-Type Live-Cell Only, Birth-Death Process

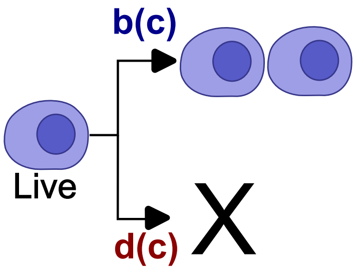

- $b$ - birth rate
- $d$ - death rate

(a) Single Dose (no concentration-response) - `model_library/1type-livecell/singledose/`

i. No Error, No Mixed Effects `./birthdeath.stan`

ii. Error, No Mixed Effects `./birthdeath_error.stan`

iii. No Error, Mixed Effects `./birthdeath_mixedeffects.stan`

iv. Error, Mixed Effects `./birthdeath_error_mixedeffects.stan`

(b) 4-parameter logistic - `model_library/1type-livecell/logistic/`

$$b(c) = bi + \frac{db}{\exp(bh(c - b50))}$$
$$d(c) = bd + dd - \frac{dd}{\exp(dh(c - d50))}$$

i. No Error, No Mixed Effects `./birthdeath.stan`

ii. Error, No Mixed Effects `./birthdeath_error.stan`

iii. No Error, Mixed Effects `./birthdeath_mixedeffects.stan`

iv. Error, Mixed Effects `./birthdeath_error_mixedeffects.stan`

(c) 2-drug Bliss combination - `model_library/1type-livecell/combination/`

$$b(c1, c2) = bi + \frac{db}{((1 + \exp(bh_1(c1 - b50_1)))(1 + \exp(bh_2(c2 - b50_2))))}$$
$$d(c1, c2) = d0 + dd - \frac{dd}{((1 + \exp(dh_1(c1 - d50_1)))(1 + \exp(dh_2(c2 - d50_2))))};$$

i. No Error, No Mixed Effects `./birthdeath.stan`

ii. Error, No Mixed Effects `./birthdeath_error.stan`

iii. No Error, Mixed Effects `./birthdeath_mixedeffects.stan`

iv. Error, Mixed Effects `./birthdeath_error_mixedeffects.stan`

(d) Gaussian Process - `model_library/1type-livecell/gaussianprocess/`

i. No Error, No Mixed Effects `./birthdeath.stan`

ii. Error, No Mixed Effects `./birthdeath_error.stan`

iii. No Error, Mixed Effects `./birthdeath_mixedeffects.stan`

iv. Error, Mixed Effects `./birthdeath_error_mixedeffects.stan`

2. 2-Type Live and Dead Cell Only, Birth-Death-Clearance Process

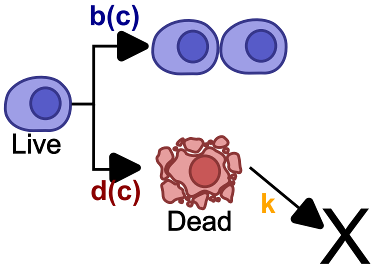

- b - birth rate
- d - death rate
- k - clearance rate

(a) Single Dose (no concentration-response) - `model_library/2type-livedead/singledose/`

i. No Error, No Mixed Effects `./birthdeathclearance.stan`

ii. Error, No Mixed Effects `./birthdeathclearance_error.stan`

iii. No Error, Mixed Effects `./birthdeathclearance_mixedeffects.stan`

iv. Error, Mixed Effects `./birthdeathclearance_error_mixedeffects.stan`

(b) 4-parameter logistic - `model_library/2type-livedead/logistic/`

$$b(c) = bi + \frac{db}{\exp(bh(c - b50))}$$

$$d(c) = d0 + dd - \frac{dd}{\exp(dh(c - d50))}$$

$$k = \text{constant}$$

i. No Error, No Mixed Effects `./birthdeathclearance.stan`

ii. Error, No Mixed Effects `./birthdeathclearance_error.stan`

iii. No Error, Mixed Effects `./birthdeathclearance_mixedeffects.stan`

iv. Error, Mixed Effects `./birthdeathclearance_error_mixedeffects.stan`

(c) Gaussian Process - `model_library/2type-livedead/gaussianprocess/`

i. No Error, No Mixed Effects `./birthdeathclearance.stan`

ii. Error, No Mixed Effects `./birthdeathclearance_error.stan`

iii. No Error, Mixed Effects `./birthdeathclearance_mixedeffects.stan`

iv. Error, Mixed Effects `./birthdeathclearance_error_mixedeffects.stan`

3. 2-Type Live Cell, Birth-Death-Mutation Process

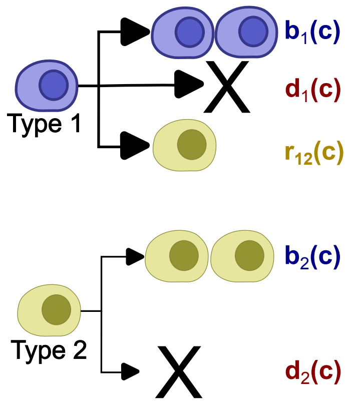

- b1 - type 1 birth rate
- d1 - type 1 death rate
- r12 - type 1 to type 2 rate
- b2 - type 2 birth rate
- d2 - type 2 death rate

(a) Single Dose (no concentration-response)

*Note: The single dose can be written as part of the k-Type Single Dose model so it is included there.*

(b) 4-parameter logistic - `model_library/2type-birthdeathmutate/logistic/`

$$b_1(c) = bi_1 + \frac{db_1}{\exp(bh_1(c - b50_1))}$$
$$d_1(c) = d0_1 + dd_1 - \frac{dd_1}{\exp(dh_1(c - d50_1))}$$
$$b_2(c) = bi_2 + \frac{db_2}{\exp(bh_2(c - b50_2))}$$
$$d_2(c) = d0_2 + dd_2 - \frac{dd_2}{\exp(dh_2(c - d50_2))}$$
$$r_12(c) = r12_0 + r12_d - \frac{r12_d}{\exp(r12_h(c - r12_50))}$$

i. No Error, No Mixed Effects `./birthdeathmutate.stan`

ii. Error, No Mixed Effects `./birthdeathmutate_error.stan`

iii. No Error, Mixed Effects `./birthdeathmutate_mixedeffects.stan`

iv. Error, Mixed Effects `./birthdeathmutate_error_mixedeffects.stan`

4. 2-Type Cell Cycle Model

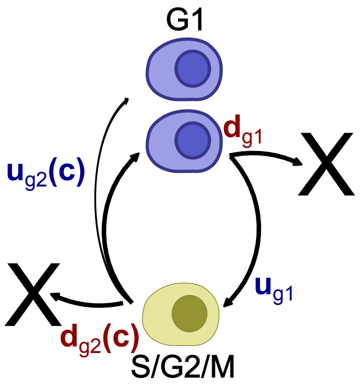

- ug1 - G1 to S/G2/M rate
- dg1 - G1 death rate

- ug2 - S/G2/M rate to 2x G1 rate
- dg2 - S/G2/M death rate

(a) Single Dose (no concentration-response)

*Note: The single dose can be written as part of the k-Type Single Dose model so it is included there.*

(b) 4-parameter logistic - `model_library/2type-cellcycle/logistic/`

$$u_g1(c) = \text{constant}$$
$$d_g1(c) = \text{constant}$$
$$u_g2(c) = ui_g2 + \frac{du_g2}{\exp(uh_g2(c - u50_g2))}$$
$$d_g2(c) = d0_g2 + dd_g2 - \frac{dd_g2}{\exp(dh_g2(c - d50_g2))}$$

i. No Error, No Mixed Effects `./cellcycle-2phase.stan`

ii. Error, No Mixed Effects `./cellcycle-2phase_error.stan`

5. 3-Type Live, Apoptotic, Dead model

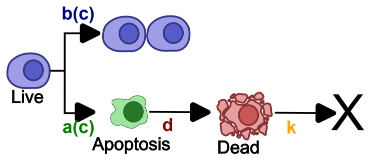

- b - birth rate (splitting)
- a - apoptosis rate (live to apoptotic)
- d - death rate (apoptotic to dead)
- k - clearance rate (dead to not tracked)

(a) Single Dose (no concentration-response)

*Note: The single dose can be written as part of the k-Type Single Dose model so it is included there.*

(b) 4-parameter logistic - `model_library/3type-liveapoptoticdead/logistic/`

$$
\begin{aligned}
b(c) &= bi + \frac{db}{\exp(bh(c - b50))} \\
a(c) &= a0 + da - \frac{da}{\exp(ah(c - a50))} \\
d &= \text{constant} \\
k &= \text{constant}
\end{aligned}
$$

i. No Error, No Mixed Effects `./birthapoptosisdeath.stan`

ii. Error, No Mixed Effects `./birthapoptosisdeath_error.stan`

6. 4-Type Cell Cycle Model

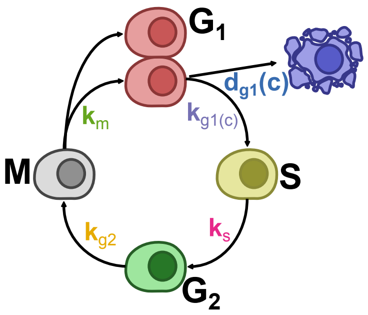

- k_G1 - G1 to S
- k_S - S to G2
- k_G2 - G2 to M
- k_M - M to 2X G1
- d_G1 - G1 death

(a) Single Dose (no concentration-response)

*Note: The single dose can be written as part of the k-Type Single Dose model so it is included there.*

(b) 4-parameter logistic - `model_library/4type-fucci/logistic/`

$$k_g1(c) = kG1_i + \frac{dkG1}{\exp(kG1_h(c - kG1_{50}))}$$
$$d_g1(c) = dG1_0 + ddG1 - \frac{ddG1}{\exp(dG1_h(c - dG1_{50}))}$$
$$k_s = \text{constant}$$
$$k_G2 = \text{constant}$$
$$k_M = \text{constant}$$

i. No Error, No Mixed Effects `./fucci_G1_doseresponse.stan`

7. k-Type Custom Model

(a) Single Dose (no concentration-response) - `model_library/ktype/singledose`

i. No Error, No Mixed Effects `./ktype.stan`

ii. Error, No Mixed Effects `./ktype_error.stan`

(b) Template for customizing a model and dose-response
`model_library/ktype/multidose-template`

#### 4.2 Custom Modeling

BESTDR can create custom models where you can specify the mechanistic model that describes how each type reproduces, dies, transitions, etc. We specify each event as a transition where the parent, offspring, and concentration-response are all explicitly stated. Additionally we can include any parameter constraints, prior information, hierarchical variables and priors, as well as whether or not to include an error term. A couple steps and functions are required to go from model specification to creating the Stan file that can be compiled. We include helper functions to add data as well.

Additionally, the code can be modified from template files to fit specific needs. The Stan code has regions where the user needs to insert

- prior parameter names (under `data` block)

- dose-response parameters (under `parameters` block)
- dose-response functions (under `transformed parameters`
- prior distributions (under `model block`)
